# Early development of direction selectivity in higher visual cortex

**DOI:** 10.1101/2025.05.15.654265

**Authors:** Dallas C. Khamiss, Augusto A. Lempel, Brandon R. Nanfito, Kristina J. Nielsen

**Affiliations:** Solomon H. Snyder Department of Neuroscience, Johns Hopkins University School of Medicine, Baltimore, MD 21205, USA; Zanvyl Krieger Mind/Brain Institute, Johns Hopkins University, Baltimore, MD 21218, USA

## Abstract

A fundamental aspect of visual motion processing is the computation of motion direction. In ferrets, as in primates, selectivity for motion direction is found both in early cortical stages like the primary visual cortex (V1) and in higher visual areas like the middle temporal (MT) area in primates and the postero-medial lateral suprasylvian (PMLS) area in ferrets. Little is known about how this critical tuning function develops in higher visual cortex. Here, by studying the development of the ferret’s motion pathway, we first reveal the surprising finding that direction selectivity develops earlier in PMLS than in V1, contrary to the areas’ hierarchical positions. Our data, collected in animals of either sex, furthermore show that while direction selectivity is sensitive to visual experience in both areas, the sensitivity profile differs between them: Presentation of drifting gratings, containing the full complement of spatial and temporal cues generated by visual motion, can promote direction selectivity development in V1 and PMLS. In contrast, flashing stationary stimuli, which lack the spatial displacement of moving stimuli and only contain temporal changes, induce direction selectivity only in PMLS, not V1. Collectively our findings reveal significant deviations in PMLS development from that in V1, which will be important to account for in models of motion pathway development and of the developmental disorders that affect this pathway. The complex pattern of relative PMLS and V1 development also highlights the need to address interactions between areas in developmental research.

**Significance Statement:** While the development of early stages of visual cortex up to primary visual cortex (V1) has received much attention throughout the years, significantly less is known about that of higher visual cortex both on its own as well as in relationship to V1. Here, we focus on a core motion function, direction selectivity, to systematically characterize the coordinated development of multiple stages of the visual motion pathway in ferrets. Crucially, our data show that this coordinated development is surprising complex, and that the developmental status of the higher areas cannot be predicted based on that in lower areas. These findings may also provide clues why motion vision is particularly vulnerable to developmental disorders.

## Introduction

Encoding of motion information is a core visual function with high behavioral relevance. Not surprisingly, circuits computing motion have been found in a wide range of species, from flies (Borst and Groschner, 2023), fish (Bollmann, 2019) to mammals, including rodents (Wei and Feller, 2011; Vaney et al., 2012; Niell and Scanziani, 2021) and primates (e.g., Born and Bradley, 2005; Orban, 2008; Chaplin et al., 2018). A particular challenge for motion computations is that they require precise processing of spatial and temporal information, which might be one reason why deficits in motion processing occur in a number of developmental disorders (Atkinson, 2017). Both their behavioral relevance and apparent vulnerability make the development of circuits supporting motion computations an important topic for investigation.

One of the most fundamental motion computations is the extraction of motion direction, which in primates and carnivores predominately occurs in cortex. Importantly, it occurs not just in early stages like the primary visual cortex (V1) (Hubel and Wiesel, 1968; De Valois et al., 1982), but also in higher visual areas. One central higher motion area in primates is the middle temporal area (MT) (Born and Bradley, 2005; Orban, 2008), which receives direct inputs from V1. In addition to exhibiting strong direction selectivity (DS), a hallmark MT function is the computation of global motion signals. In carnivores, the postero-medial lateral suprasylvian sulcus area (PMLS or PSS in ferrets, see Methods) is thought to play a similar role. Similarities between ferret PMLS and MT includedirect inputs from V1 (Jarosiewicz et al., 2012; Khalil et al., 2018), and a high degree of DS (Philipp et al., 2006; Lempel and Nielsen, 2019). Importantly, we have recently shown that PMLS neurons signal global motion directions (Lempel and Nielsen, 2019).

Research on DS development at the neural level has so far almost exclusively focused on V1 (e.g. Li et al., 2006; Clemens et al., 2012; Smith et al., 2015; Chang and Fitzpatrick, 2022). Very little is currently known about its emergence in crucial higher motion areas like MT (Movshon et al., 2004) or PMLS (Price et al., 1988), and even less about how it compares to V1. Here, we address this gap in knowledge regarding the development of a key motion computation by systematically following the maturation of DS in ferret PMLS, and directly comparing it to V1.

In addition to the time course of maturation, another important aspect of visual development is the degree to which it is sensitive to visual information – both its general presence, as well as its specific content. Indeed, providing or withholding certain types of visual information influences DS development in ferret V1 (Li et al., 2006, 2008; Van Hooser et al., 2012; Ritter et al., 2017; Roy et al., 2020). Altered visual experience also affects DS in cat PMLS and monkey MT (Spear et al., 1985; El-Shamayleh et al., 2010; Grootel et al., 2024), and early motion integration development in ferret PMLS shows influences of visual experience (Lempel and Nielsen, 2021). Visual experience therefore appears to play a role in circuit development in higher motion cortex in addition to V1. However, all of these studies investigated a single area at a time, and a systematic comparison of the effects of different types of visual experience across the visual hierarchy is lacking. Here, we directly compare the impact of different visual cues on V1 and PMLS development, and use it as a sensitive probe for the developmental processes shaping DS in V1 and PMLS.

In general, our results show a surprising degree of divergence between the two areas, with PMLS DS development occurring earlier, and under the influence of different types of visual information, than in V1. These results strongly suggest that lower stages do not always set the pace for the development of the higher areas they provide input to, and highlight the need to consider multiple areas and their interactions in models of motion pathway development.

## Materials and Methods

### Animal preparation, surgery and recording location

All procedures adhered to the guidelines of the National Institute of Health and were approved by the Animal Care and Use Committee at Johns Hopkins University. Experiments were performed in male and female ferrets (*Mustela putoris furo*) aged 29 -52 days. Every animal contributed to a single time point only. Animals used in the training experiments were 28 -33 days old; none of these animals had naturally opened their eyes at the start of the experiment. Recordings were performed in anesthetized ferrets following established procedures described previously (Lempel and Nielsen, 2019). Briefly, animals were anesthetized using isoflurane (during surgery: 1.5-3%, during recordings: 0.5-2%), and paralyzed using pancuronium bromide (0.15 mg/kg/hr) to prevent eye movements. A range of vital parameters (heart rate, SpO2, EtCO2, EEG) were monitored continuously to ensure adequate anesthetic depth during the experiments. Neosynephrine and atropine were applied to the eyes to retract the nictitating membranes and dilate the pupils. Eyes were then covered with contact lenses.

To record neural signals, craniotomies were made over V1 and PMLS. When recording from both areas simultaneously, we targeted related visual field positions in both areas. PMLS is located in the posterior bank and fundus of the suprasylvian sulcus, close to its medial end (Fig. 1A). In the literature, this area has been referred to by a number of different names, all of which are based on its anatomical location, but with varying degrees of specificity. The three names generally in use (with example publications listed) are SSY for suprasylvian sulcus (Bizley et al., 2007; Khalil and Levitt, 2014), PSS for posterior suprasylvian sulcus (Philipp et al., 2006; Lempel and Nielsen, 2019) and PMLS for posterio-medial suprasylvian sulcus (Homman-Ludiye et al., 2010). PMLS, as the most specific label, differentiates the area from other areas located in the suprasylvian sulcus more precisely and will be used here.

**Figure 1:**
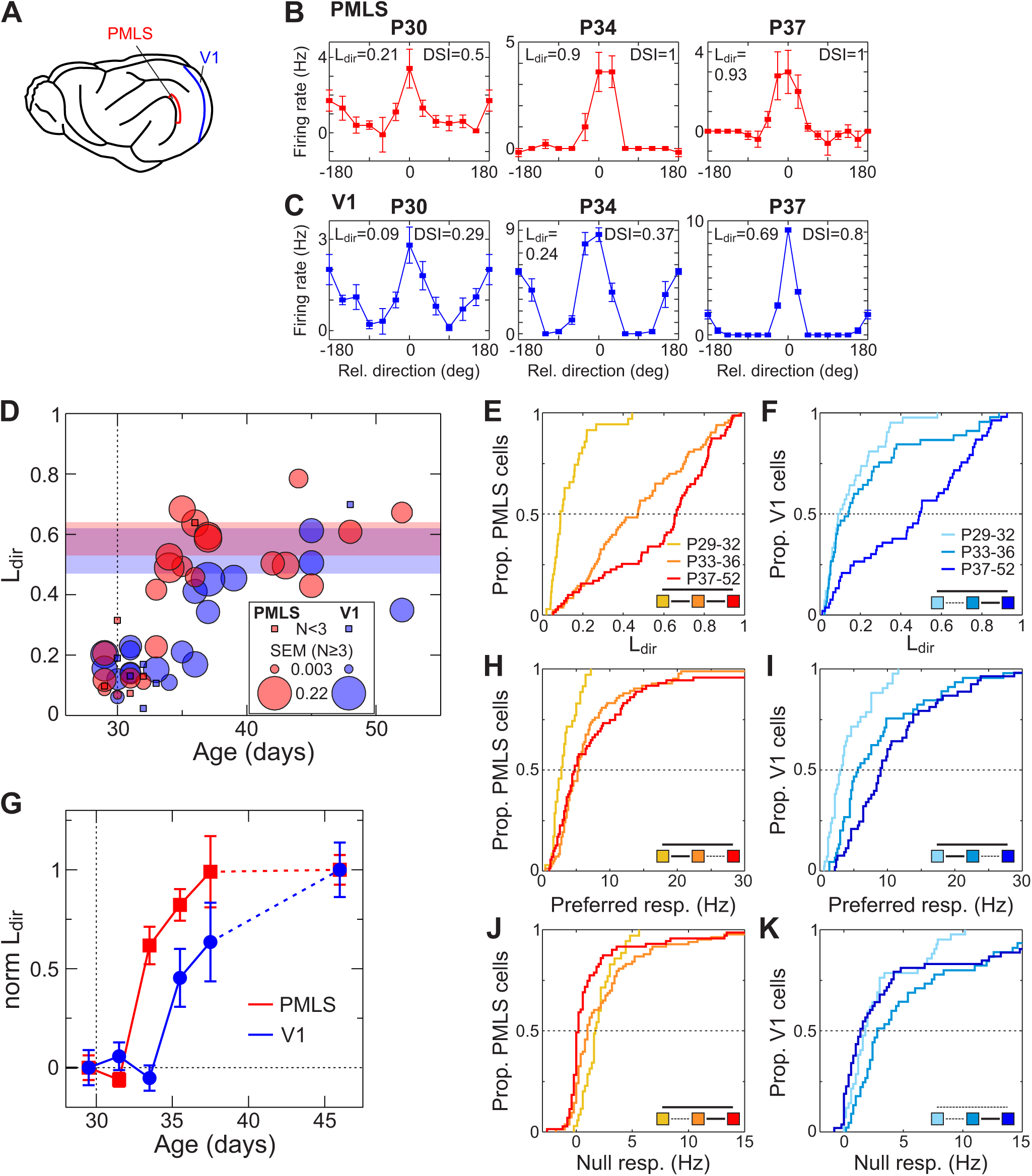
Development of DS in PMLS and V1. (A) Side view of the ferret brain showing the locations of V1 (or area 17) and PMLS (or PSS). (B) Responses of 3 PMLS neurons to drifting gratings, recorded at different ages (indicated above each plot). Grating direction is plotted relative to each neuron’s preferred direction. Measures for DS (L_dir_ and DSI) are given for each neuron. (C) Responses of 3 V1 neurons, following the same format as (B). (D) Development of L_dir_ on a per-animal basis for V1 and PMLS. Each circle indicates the mean L_dir_ across all visually responses SU in an animal, as a function of the animal’s age. Circle size corresponds to the SEM for the mean for animals in which 3 or more neurons were recorded. Means based on less than 3 neurons are shown with squares. Note that the SEM is plotted using a separate scale from the y-axis to maintain visibility of all data points. The legend shows the scale for the SEM. The colored bands indicate the mean +/- 1 SEM for animals 40 days and older. The dashed line at P30 indicates average age of eye opening in ferrets. (E) Cumulative distribution of SU L_dir_ values recorded in PMLS for 3 age groups. The legend in the bottom right corner indicates the outcome of rank-sum tests between the different groups (dashed line: p>.01; solid line: p<.01). (F) Cumulative distribution of SU L_dir_ values for V1. Same format as (E). (G) Normalized L_dir_ as a function of age for V1 and PMLS. SU L_dir_ data were divided into 2-day age bins from P29-38, and one adult bin from P40-52. Data were normalized independently for each area so that the average normalized L_dir_ in the P29-30 group equals zero, and the average L_dir_ for P40-52 equals 1. The normalized data were then averaged within each age bin and are plotted as a function of the middle of each age bin. Error bars: Standard error of mean (SEM). (H, I) SU firing rates evoked by the preferred direction for PMLS and V1. Same format as (E, F). (J, K) Distribution of responses evoked by the null direction for V1 and PMLS, same format as (E, F).

### Extracellular recordings and extraction of neural responses

Neural responses were recorded using multichannel silicon probes obtained from the Masmanidis lab at UCLA (Du et al., 2011), plated with gold to reach impedances around 150-300 kΩ. Signals from one or two probes were recorded with a RHD2000 amplifier (intan Technologies) using sampling rates of 20-30 kHz. Data was filtered offline between 250 Hz and 5kHz. Based on these filtered signals, two types of neural responses were extracted. Single units (SU) were isolated using a custom-written processing pipeline in Matlab (MathWorks) for manual spike sorting (available on the nielsenlab github repository). In this pipeline, detection thresholds were first set manually for each recording channel based on noise levels, and used to extract spikes for each channel. Spikes occurring within +/-0.2 ms of each other on channels with distances ≤50 μm were assigned to the channel with the largest energy and eliminated from other channels. Spikes were then manually clustered using multiple waveform characteristics like trough amplitude, energy, or trough-peak amplitude, as well as recording position on the probe (determined as center-of-mass of the spike energy or trough amplitude computed across neighboring probe channels). Quality of isolation was confirmed by analyzing inter-spike intervals and eliminating any units with intervals below 1.2 ms. Multi-unit (MU) signals were extracted by first applying an automatically computed detection threshold to every channel. The threshold was set to 4 times the standard deviation of each channel’s bandpass-filtered signal, using the median absolute deviation scaled for a normal distribution to estimate the standard deviation. Again, events detected within +/-0.2 ms on channels within 50 μm were assigned only to the channel with the largest energy. No further spike sorting was performed on the MU signal.

### Visual stimuli and experiment design

#### Visual stimulus generation

Stimuli were generated using the Psychophysics Toolbox extension to Matlab (Brainard, 1997; Pelli, 1997) and displayed on a 23 inch LCD monitor with a refresh rate of 120 Hz, placed 20 – 35 cm in front of the ferret. The monitor was gamma corrected using a SpectraScan 655 (PhotoResearch).

#### Stimuli for selectivity measurements

Except for the stimulation blocks of the training paradigm, all data were collected by presenting gratings drifting in 12 or 16 directions. In addition to the different directions, each experiment also contained a blank condition. All conditions were shown in a pseudo-random sequence with 5 repetitions per condition. Gratings were shown full-field (about 65 x 50 deg), with the exception of 3 experiments that also presented an additional smaller size. For those experiments, only the full-field data was analyzed. Gratings were square-wave gratings shown at 100% contrast for all but two experiments in adult animals (at P45 and P48), which used sine-wave gratings instead.

Grating spatial and temporal frequency were optimized to drive robust responses at each recording site. In cases in which PMLS and V1 data were collected simultaneously, spatial and temporal frequency values were chosen based on the collective responses in both areas. Importantly, this also means that they were the same for both areas in these simultaneous recording experiments. Spatial and temporal frequency values are plotted for all experiments as a function of recording area and age in Fig. S1. In general, values for spatial frequency ranged from 0.04 – 0.1 cyc/deg for PMLS and 0.05 – 0.1 cyc/deg in V1 (but it should be noted that 0.04 cyc/deg was only used in a single PMLS experiment). Temporal frequency ranged from 2 – 6 cyc/s for PMLS and 1 -6 cyc/s for V1 (with a single V1 experiment using 1 cyc/s). Each grating presentation was preceded and followed by presentation of a gray screen of equal luminance. Stimuli were shown for 1 s, with inter-stimulus intervals (ISI) of 3 -10 s. Longer intervals were needed for the youngest animals in which adaptation to the stimulus is more pronounced (see Fig. S1 for a plot of ISI values across all experiments). Again, ISI values were set to optimal values for both V1 and PMLS recording sites collectively in simultaneous recording sessions. Across all experiments, ISI values for PMLS ranged from 4 to 10 s, and for V1 from 3 to 10 s (with a single experiment using a 3 s ITI).

#### Stimuli during training paradigm

For the animals exposed to drifting or flashing stimuli, we presented full-field, square-wave gratings during the training blocks. Stimuli were shown for 5 s, with a 10 s ISI. Spatial frequency was again adjusted based on responses in an individual animal, and ranged between 0.05 – 0.08 cyc/deg for both groups. For the drifting gratings, temporal frequency ranged between 3 – 6 cyc/deg. Drifting gratings were always shown at 100% contrast. For the flashing gratings, we modulated contrast between 0 and 100% (no contrast reversal) using a sinusoidal function with a temporal frequency of 3 Hz.

### Data set size and inclusion criteria

For the analysis of SU responses, neural responses to every stimulus were calculated as the firing rate during the stimulus presentation period (1 s long) minus the firing rate during the last second of the pre-stimulus period (adjusting the start of the stimulus window to account for developmental changes in latency does not impact the results). SU needed to pass two criteria to be included in the analyses. First, we computed an ANOVA across the responses in all stimulus conditions, including the blank. Only SU with p<.05 for the ANOVA were included. Additionally, the average firing rate for the best condition had to exceed 0 Hz.

Experiments included in the SU analysis were generally recorded early in the recording session in an animal, so that stimulus exposure prior to data collection was kept to a minimum. More specifically, 24 out of 28 V1 experiments were preceded by less than 10 min of cumulative stimulus exposure (mean cumulative stimulus exposure prior to data collection in these experiments: 1.3 min). 3 of the remaining 4 experiments occurred after 47 min of stimulus exposure, and one experiment after 3 h of stimulus presentation. All of these 4 experiments were performed in animals P39 and older. For PMLS, 25 of 30 experiments occurred after less than 10 min of stimulus exposure (mean cumulative stimulus exposure for these experiments: 1.4 min). The remaining 5 experiments (1 at P30, the rest at >P42) were preceded by 40 – 48 min of stimulus exposure. Thus, there is in general little opportunity for prior stimulus exposure to impact the tuning levels reported here. For V1, data collection after longer stimulus exposure only occurred in mature animals. Stimulus exposure duration therefore does not affect any of the conclusions regarding development during the week after eye opening. For PMLS, the single data set after longer stimulus exposure collected at P30 contributed 3 neurons (out of 35) to the respective age group; all other cases again are mature animals. Excluding the P30 data from the analysis does not impact any of the conclusions.

For the analysis of response latency using MU data, we used the same approach to test for visual responsiveness (ANOVA for the baseline-corrected firing rates with p<.05, firing rate for best stimulus > 0 Hz). Additionally, the data set was limited to simultaneous recordings from V1 and PMLS. The resulting MU data set includes the same recordings used for SU analysis of DS development, but contains additional data in which no SU could be isolated or in which no SU passed the criteria.

MU data was also used to analyze the effects of the training paradigm. In this case, the long duration of the experiments generally made responses more variable, which required an adjustment in the inclusion criteria. For these data, we tested for visual responsiveness by comparing the stimulus-evoked firing rates (not baseline corrected) to the spontaneous activity during the 1 s-interval preceding each stimulus presentation using a rank-sum test (blanks were excluded from this analysis). MU with p<.05 and with responses to the preferred stimulus exceeding 0 Hz were included in subsequent analyses. Further, data were restricted to experiments in which visually responsive sites could be identified before and after training. This meant that for a few animals only PMLS data (1 animal in the flashing and gray cohort each) or V1 data (1 animal in the gray cohort) were available. These cases were eliminated when comparing training effect sizes between V1 and PMLS (e.g., Fig. 4H). Finally, no responsiveness criterion was applied to the MU collected during the training blocks. The resulting number of animals and neurons for normal development and training experiments are listed in Table 1.

**Table 1:**
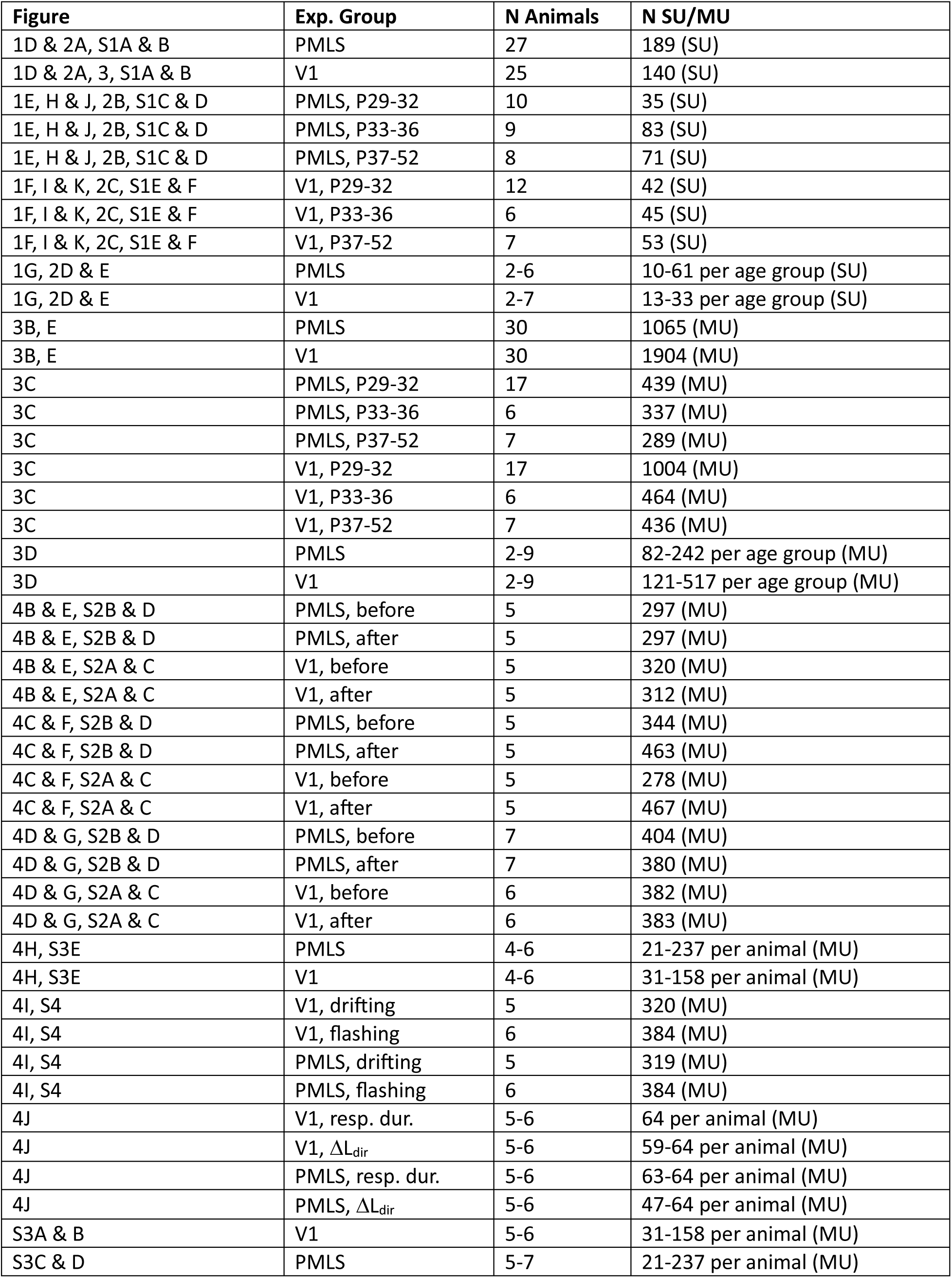
Data set sizes for all figures and analyses.

| Figure | Exp. Group | N Animals | N SU/MU |
| --- | --- | --- | --- |
| 1D & 2A, S1A & B | PMLS | 27 | 189 (SU) |
| 1D & 2A, 3, S1A & B | V1 | 25 | 140 (SU) |
| 1E, H & J, 2B, S1C & D | PMLS, P29-32 | 10 | 35 (SU) |
| 1E, H & J, 2B, S1C & D | PMLS, P33-36 | 9 | 83 (SU) |
| 1E, H & J, 2B, S1C & D | PMLS, P37-52 | 8 | 71 (SU) |
| 1F, I & K, 2C, S1E & F | V1, P29-32 | 12 | 42 (SU) |
| 1F, I & K, 2C, S1E & F | V1, P33-36 | 6 | 45 (SU) |
| 1F, I & K, 2C, S1E & F | V1, P37-52 | 7 | 53 (SU) |
| 1G, 2D & E | PMLS | 2-6 | 10-61 per age group (SU) |
| 1G, 2D & E | V1 | 2-7 | 13-33 per age group (SU) |
| 3B, E | PMLS | 30 | 1065 (MU) |
| 3B, E | V1 | 30 | 1904 (MU) |
| 3C | PMLS, P29-32 | 17 | 439 (MU) |
| 3C | PMLS, P33-36 | 6 | 337 (MU) |
| 3C | PMLS, P37-52 | 7 | 289 (MU) |
| 3C | V1, P29-32 | 17 | 1004 (MU) |
| 3C | V1, P33-36 | 6 | 464 (MU) |
| 3C | V1, P37-52 | 7 | 436 (MU) |
| 3D | PMLS | 2-9 | 82-242 per age group (MU) |
| 3D | V1 | 2-9 | 121-517 per age group (MU) |
| 4B & E, S2B & D | PMLS, before | 5 | 297 (MU) |
| 4B & E, S2B & D | PMLS, after | 5 | 297 (MU) |
| 4B & E, S2A & C | V1, before | 5 | 320 (MU) |
| 4B & E, S2A & C | V1, after | 5 | 312 (MU) |
| 4C & F, S2B & D | PMLS, before | 5 | 344 (MU) |
| 4C & F, S2B & D | PMLS, after | 5 | 463 (MU) |
| 4C & F, S2A & C | V1, before | 5 | 278 (MU) |
| 4C & F, S2A & C | V1, after | 5 | 467 (MU) |
| 4D & G, S2B & D | PMLS, before | 7 | 404 (MU) |
| 4D & G, S2B & D | PMLS, after | 7 | 380 (MU) |
| 4D & G, S2A & C | V1, before | 6 | 382 (MU) |
| 4D & G, S2A & C | V1, after | 6 | 383 (MU) |
| 4H, S3E | PMLS | 4-6 | 21-237 per animal (MU) |
| 4H, S3E | V1 | 4-6 | 31-158 per animal (MU) |
| 4I, S4 | V1, drifting | 5 | 320 (MU) |
| 4I, S4 | V1, flashing | 6 | 384 (MU) |
| 4I, S4 | PMLS, drifting | 5 | 319 (MU) |
| 4I, S4 | PMLS, flashing | 6 | 384 (MU) |
| 4J | V1, resp. dur. | 5-6 | 64 per animal (MU) |
| 4J | V1, $\Delta L_{dir}$ | 5-6 | 59-64 per animal (MU) |
| 4J | PMLS, resp. dur. | 5-6 | 63-64 per animal (MU) |
| 4J | PMLS, $\Delta L_{dir}$ | 5-6 | 47-64 per animal (MU) |
| S3A & B | V1 | 5-6 | 31-158 per animal (MU) |
| S3C & D | PMLS | 5-7 | 21-237 per animal (MU) |

### Measurements of direction and orientation selectivity

DS was quantified using 2 metrics, L_dir_ and DSI. L_dir_ was computed as (Mazurek et al., 2014)

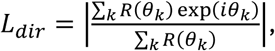

where R(θ_k_) represents the response to angle θ_k_. DSI was computed as

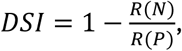

where R(P) is the response to the preferred and R(N) the response to the null direction. Matching metrics were computed for orientation selectivity (OS):

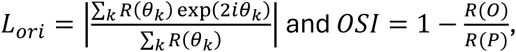

with R(O) the response to the orthogonal direction.

### Normalized DS and OS metrics

To compare development of DS and OS across areas without the effects of different selectivity levels in adults, normalized L_dir_ and L_ori_ values were computed. For this analysis, SU were first binned into 6 age windows, P29-30, P31-32, P33-34, P35-36, P37-38, P40-52. We then computed the average L_dir_ value for the first and last bin. All L_dir_ values were then normalized by computing

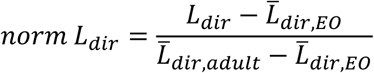

where 𝐿̅_𝑑𝑖𝑟,𝐸𝑂_ refers to the average L_dir_ value in the first age group around eye opening, and 𝐿̅_𝑑𝑖𝑟,𝑎𝑑𝑢𝑙𝑡_ to the average value in the last, adult group. The same analysis was repeated to compute normalized L_ori_ values.

### Changepoint analysis

We used Matlab’s changepoint detection function (findchangepts) to determine the onset of DS or OS maturation. Changepoints correspond to points (here, ages) with abrupt changes in the mean of the input data. The findchangepts function identifies them by partitioning the input data into two (or more) segments, divided at the changepoint, so that the sum of the squared error of each segment from its local mean is minimized. To compute changepoints, we binned the SU data from animals aged P29 to P38 into 5 two-day age windows, and then computed the mean L_dir_ or L_ori_ per age window and area. Confidence intervals for changepoints were computed using bootstrapping (bootstrp, Matlab) to resample (with replacement) the SU data in each age window, followed by computing changepoints based on the means of the resampled data, repeated 2000 times.

### Measurements of latency

Latency was computed for each MU site based on the responses to its best condition. Latencies were computed using cumulative sum analysis (Ellaway, 1978) using similar statistical criteria as used previously for primate MT neurons (Raiguel et al., 1999). First, responses were binned into 10 ms bins to compute the peristimulus time histogram (PSTH). Second, mean and standard deviation of the baseline rate were computed for the 1 s baseline preceding stimulus onset, and the baseline mean was subtracted from the PSTH before computing the cumulative sum of the PSTH. The latency was then taken as the first bin after stimulus onset in which the cumulative PSTH was larger than two standard deviations (of the baseline rate), followed by 2 increasing bins. For the analysis of latency development across two-day age windows, data from V1 and PMLS were binned into the same 6 age windows used for the normalized DS and OS metric described above.

### Analysis of MU responses in training blocks

To analyze response patterns during the training blocks, responses were computed during the full 5 s of stimulus presentation. To compute average rate, the firing rate was computed for each 5 s stimulus presentation period, baseline corrected by the firing rate in the preceding 2 s baseline period, and averaged across all stimulus presentation periods. To compute peak rate and response duration, we first computed the PSTH across all stimulus presentation periods using 10 ms bins. The peak rate was set to the maximum of the PSTH, baseline corrected as before. Duration was quantified as the number of PSTH bins during the stimulus presentation exceeding the baseline rate by 2 standard deviations, converted to seconds. Finally, all metrics were averaged across all blocks for each channel. This assumes that there are no significant movements of the probe throughout the experiment. Treating each block as an independent measurement (no averaging across blocks) does not affect the results.

### Statistics

For all normal development data, we first used an ANOVA or Kruskall-Wallis test to establish high-level differences between groups or areas (see Tables 2-13). Post-hoc tests (where justified) were then performed using Wilcoxon rank-sum tests. The outcome of all statistical tests is listed in Tables 2-13. For the training data, we performed two different kinds of analyses. First, we compared before and after training data, combined across all animals, for each area using a rank-sum test. Second, we fit the L_dir_ MU data with a linear mixed-effects (LME) model (lmefit, Matlab), with training phase (before/after exposure to the stimulus) as fixed effect and per-animal random intercepts and slopes. We compared this model against one omitting the per-animal random slopes; in most instances, the model including random slopes outperformed the one without (based on likelihood-ratio comparisons), so for consistency, we used the model with random slopes in all cases. Models were fit using maximum likelihood estimation, and the significance of the fixed effect was assessed using an ANOVA. Effect size per animal was computed as the sum of the fixed effect slope and the random, per-animal slope.

**Table 2:** Statistics for Figs. 1E, F, H-K.

| Metric | Group | Test | PMLS p value | PMLS other | V1 p value | V1 other |
| --- | --- | --- | --- | --- | --- | --- |
| L<sub>dir</sub> | All age groups | Kruskall-Wallis | <0.001 | X(2)=64.9 | <0.001 | X(2)=30.2 |
| L<sub>dir</sub> | P29-32 vs P33-36 | Rank sum | <0.001 |  | 0.3 |  |
| L<sub>dir</sub> | P29-32 vs P37-52 | Rank sum | <0.001 |  | <0.001 |  |
| L<sub>dir</sub> | P33-36 vs P37-52 | Rank sum | 0.006 |  | <0.001 |  |
| Pref. | All age groups | Kruskall-Wallis | <0.001 | X(2)=20.6 | <0.001 | X(2)=34.3 |
| Pref. | P29-32 vs P33-36 | Rank sum | <0.001 |  | <0.001 |  |
| Pref. | P29-32 vs P37-52 | Rank sum | <0.001 |  | <0.001 |  |
| Pref. | P33-36 vs P37-52 | Rank sum | 0.9 |  | 0.03 |  |
| Null | All age groups | Kruskall-Wallis | <0.001 | X(2)=22.1 | 0.008 | X(2)=9.6 |
| Null | P29-32 vs P33-36 | Rank sum | 0.2 |  | 0.01 |  |
| Null | P29-32 vs P37-52 | Rank sum | <0.001 |  | 0.6 |  |
| Null | P33-36 vs P37-52 | Rank sum | <0.001 |  | 0.006 |  |

## Results

### After eye opening, DS develops faster in PMLS than V1

DS develops in ferret V1 in the week after eye opening, and matures around postnatal day (P) 37 (Li et al., 2006; Clemens et al., 2012). While we have previously shown that PMLS DS is mature at the same age (Lempel and Nielsen, 2021), the time course of maturation has not yet been determined. Here, we systematically tracked PMLS DS development and compared it directly to V1. To this end, we recorded responses to drifting gratings in both areas from P29 (around eye opening) to P52 (young adult). Recordings were performed in anesthetized animals using multi-channel silicon probes, in many cases collecting data from V1 and PMLS simultaneously by recording from 2 probes. All data were collected with the eyes open, irrespective of the natural eye opening state of an animal. Fig. 1B & C show example tuning functions for well isolated SU in both areas from P30 to P37. As these plots show, we were able to find neurons with reliable visual responses at all ages, which allowed us to quantify tuning properties. The remainder of the analyses in this section will be limited to these visually responsive single neurons (determined using an ANOVA across mean baseline-corrected firing rates for all conditions including the blank with a criterion of p<.05; see Methods). The resulting data set sizes are listed in Table 1.

Especially at the youngest ages, tuning functions were often poorly fit with standard direction (or orientation) tuning functions. We therefore quantified DS by computing two different metrics. The first was L_dir_ or 1-DirCircVar, a vector-based metric (Mazurek et al., 2014) that takes the entire tuning function into account (see Methods for definition). Importantly, L_dir_ = 1 corresponds to high selectivity (responses to a single direction only), while L_dir_ = 0 indicates no selectivity (equal responses to all conditions). This metric is comparable to circular variance (CircVar), commonly used to quantify orientation tuning strength (Batschelet, 1981). The second metric used to quantify DS was the direction selectivity index (DSI), which compares only the responses between the preferred direction and the one opposing it (the null direction; see Methods). Results were generally identical between the two metrics, so only the results based on L_dir_ (which is more robust because it is based on the entire tuning curve) are discussed further (see Fig. S2 and Table 13 for the key DSI results). Both L_dir_ and DSI values are indicated for each example neuron in Fig. 1B & C, and suggest a general increase in DS with age in both areas.

To capture the time course of this DS maturation and to compare it between areas, we next analyzed DS as a function of age. As a first step, we computed the mean L_dir_ for each individual animal by averaging across all visually responsive PMLS or V1 neurons collected in that animal (results based on the median are comparable; data not shown). This makes it possible to visualize developmental trends without having to group animals by age; it also more clearly shows variability in developmental stage across animals of the same age. Fig. 1D plots these per-animal means as a function of age. The two horizontal bands correspond to the mean +/- 1 standard error (SEM) for animals P40 and older. Consistent with previous data (Li et al., 2006; Clemens et al., 2012) from ferret V1, our data show V1 DS to be immature around P30. PMLS DS levels were similarly low at this age. L_dir_ values eventually improved to mature levels in both areas, with some of the improvements occurring very rapidly, as evident by significant changes in DS between animals with a single day age difference. Unexpectedly, this sudden developmental improvement occurred earlier in PMLS than V1 – around P34 for PMLS, P36 for V1. The faster PMLS development continued, in that mature values were reached faster after the initial improvements than in V1. Judging by the overlap between the per-animal mean and the young adult range, PMLS matured around P35, but it took V1 until at least P40 to reach this point.

This difference between PMLS and V1 development was also apparent when dividing single unit L_dir_ values for each area into 3 age groups used previously for ferret V1 development (collapsing data across animals). The youngest of the 3 groups corresponded to eye opening (P29-32), the second to the phase of rapid DS development (P33-36), and the last to animals with mature DS levels (P37 and older). Fig. 1E & F plot the L_dir_ distributions for these groups for PMLS and V1. Age had a general effect on the L_dir_ distribution in both areas (Kruskal-Wallis test across all 3 groups: p<.001 for both areas; full statistics in Table 2, including the p-values for all non-significant tests). For PMLS, post-hoc rank-sum tests revealed that all three age groups deviated from each other (p<.001). In V1, on the other hand, the P29-32 & P33-35 groups were not significantly different, but both differed from the oldest group (p<.001).

It should be noted that the oldest age group here only includes ‘young adults’ up to P52 because we focused on the rapid initial phase of DS development. It is possible that DS levels in either area exhibit further changes over much longer time scales. The mean L_dir_ in PMLS in the oldest group here was 0.59 ± 0.08 (95% confidence interval (CI)). We have previously observed a mean L_dir_ of 0.56 ± 0.05 for animals aged P57 – P497 (Lempel and Nielsen, 2019), which suggests that at least PMLS L_dir_ values remain stable after the early developmental phase studied here. In V1, the mean L_dir_ in the oldest group equaled 0.46 ± 0.08, with a median value of 0.49. In our previous study in animals aged P57 – 497, the mean L_dir_ equaled 0.29 ± 0.06 in V1 (Lempel and Nielsen, 2019). This would suggest that V1 L_dir_ levels are transiently higher around P50 than in fully mature animals. Indeed, V1 DS levels have been shown to trend down from P60 to P90 (although that trend did not reach significance), with the mean DSI falling from about 0.7 to 0.5 from P60 to P90 (Popovic et al., 2018). This previous P60 DSI value is in close agreement with our data here (mean DSI for the oldest group here: 0.71 ± 0.08).

Since development is obviously dynamic and averaging across age groups cannot capture all trends, Fig. 1G uses an alternate analysis approach to show the relative DS development in V1 and PMLS in more detail. Here, L_dir_ values between P29 and 38 were binned into much smaller age windows of 2 days, combining data across animals. We also added a final bin from P40-52 (corresponding to the mature levels indicated in Fig. 1D). Since in adult animals DS is stronger in PMLS than V1, the comparison of the developmental trajectories between the areas is complicated by the fact that they reach different endpoints. To account for these endpoint differences, we chose to normalize L_dir_ values so that the average L_dir_ in the 2-day window around eye opening (P29 to 30) was set to 0, and the average L_dir_ values in the P40-52 was set to 1 (see Methods). V1 and PMLS data were normalized independently, and the normalized L_dir_ values were averaged in each age bin. These normalized time courses again show that PMLS DS development occurred earlier than in V1. PMLS L_dir_ values rapidly increased starting with the P33-34 window, while V1 L_dir_ values remained at the eye-opening level until P35-36. A two-way ANOVA with factors area and age confirmed these observations. For the ANOVA, the age windows used for normalization were excluded (since they are forced to be 0 and 1, respectively), reducing the data to P31 – 38. For these data, the two-factor ANOVA then resulted in a significant effect for age (p<.001), consistent with the general age-dependent change in DS in both areas. Importantly, a significant effect of area (p<.001), as well as an interaction between age and area trending towards significant (p=.01), support differences in DS levels between areas during this time period, which because of the normalization cannot be explained by overall higher DS levels in PMLS, but reflect developmental differences. Further post-hoc rank-sum tests indeed revealed significantly higher normalized L_dir_ values in PMLS than in V1 at P33-34 (p<.001) and P34-35 (p=.008), confirming the earlier development of DS in PMLS (see Table 3 for full test statistics). It is worth noting that the time courses in Fig. 1G also suggest that V1 DS development does not just start later, it also takes longer to complete (note the shallower slope of the V1 time course).

**Table 3:** Statistics for Figs. 1G (IA: Interaction)

| Metric | Group | Test | p value | Other |
| --- | --- | --- | --- | --- |
| Norm. L<sub>dir</sub> | V1 vs PMLS | ANOVA: area, age | Age: <0.001<br>Area: <0.001<br>IA: 0.01 | Age: F(3)=16.9<br>Area: F(1)=13.1<br>IA: F(3)=3.9 |
| Norm. L<sub>dir</sub> | V1 vs PMLS, P31-32 | Rank sum | 0.53 |  |
| Norm. L<sub>dir</sub> | V1 vs PMLS, P33-34 | Rank sum | <0.001 |  |
| Norm. L<sub>dir</sub> | V1 vs PMLS, P35-36 | Rank sum | 0.008 |  |
| Norm. L<sub>dir</sub> | V1 vs PMLS, P37-38 | Rank sum | 0.1 |  |

As a final comparison of the developmental trajectories between areas, we took advantage of the fact that L_dir_ values increased rapidly in both areas once maturation set in, but were relatively stable values before and after, to determine the onset of maturation in PMLS versus V1. To this end, we binned the (non-normalized) SU data from P29-38 into the same 2-day windows used for the analysis above, and computed the mean L_dir_ per age bin per area. The oldest group reflecting the more steady-state adult data was excluded here since the focus was on the rapid initial transition in DS levels. Data were binned to increase sample size per bin, but an analysis without binning yielded similar results. We then used Matlab’s findchangepts function to divide the average L_dir_ time course per area into two segments with locally stable means. Specifically, the algorithm determines the changepoint as the sample (here, age bin) that divides the data set into two segments such that the residual error between each segment’s data and the segment mean is minimized. A 95% CI was determined for every changepoint using bootstrapping (see Methods). For PMLS L_dir_, this approach indicated a change at P33 (CI: P33-35), while V1 L_dir_ did not change until P35 (CI: P35-37), further confirming the lag in DS development in V1 relative to PMLS.

At the most fundamental level, high DS requires strong responses to the preferred direction, and weak responses to the opposite (null) direction. To probe some of the mechanisms underlying DS development, we compared the baseline-corrected firing rates evoked by preferred and null direction at different ages. To capture the overall trends, data were divided into the same 3 age groups used for Fig. 1E & F. In both areas, the responses to the preferred stimulus became more robust with age (Fig. 1H & I, see Table 2 for statistics). Kruskall-Wallis tests comparing the 3 age groups were significant for both areas (both p<.001), with significant differences between the two youngest groups and the oldest group in both areas (rank-sum tests, all p<.001). The increase in the V1 preferred direction response is consistent with previous reports (Clemens et al., 2012); our data now show that a similar increase occurs in PMLS.

Responses to the null stimulus changed with development as well, but with differences across areas (Fig. 1J & K). In V1, null direction responses temporarily increased in the P33-36 group, a change that reached significance in comparison to the oldest group, and trended toward significance relative to the P29-32 group (Kruskall-Wallis test: p=.008; rank-sum tests P33-36 versus P37+: p=.006, P33-36 versus P29-32: p=.01). The difference between the two older groups parallel a previous observation of a drop in V1 null responses after P35 (Clemens et al., 2012). In

PMLS, on the other hand, null direction responses decreased with age, so that the oldest group had the lowest null responses (Kruskall-Wallis and post-hoc test between both younger and the oldest age group, p<.001; Table 2). This weaker response to the null direction, especially because it was paired with an increase in the preferred direction response, would support better discriminability of the two directions, and points to a possible role of null direction inhibition especially in the development of PMLS DS (see also Discussion).

Preferred and null responses can also be used to begin to address another question regarding the data. Even in adult animals, the transformation of information from V1 to PMLS results in enhanced DS in PMLS. It is possible that the same mechanisms – like integration across V1 inputs and threshold nonlinearities (Rust et al., 2006; Lempel and Nielsen, 2021) – amplify subtle direction biases present in V1 early in life, and that these biases increase in V1 during the phase of rapid PMLS DS development ahead of the major changes in DS in V1. Ultimately, measurements of the V1 input to PMLS neurons are required to fully answer this question. The data shown in Fig. 1, however, argue against this explanation: At a per-animal level, animals with mean L_dir_ levels exceeding the levels found around eye opening can be found first at P33 in PMLS, but only at P36 in V1 (Fig. 1D). Fig. 1G similarly shows that the first bin in which the normalized L_dir_ values deviate from 0 is P33-34 for PMLS and P35-36 for V1. As long as the entire tuning function is considered, our data therefore show stable tuning levels in V1 while PMLS levels are rapidly changing, inconsistent with the notion of small V1 shifts supporting the PMLS development. However, subtle shifts may be hard to detect at the level of the entire tuning function. We therefore also computed the ratio of null to preferred responses for every neuron, as a way to capture the relative strength of the null direction response independent of changes in overall firing rates (the DSI represents an alternate way to quantify the relative null direction response and yields the same results). Comparing ratios between V1 and PMLS then provides a sense of the amplification of V1 direction differences present in the PMLS output. If there were subtle changes in direction biases in V1 after eye opening, these would be reflected in decreasing ratios during this time period. Fig. S2 G-I plot the response ratios for V1 and PMLS divided into the same 3 age groups used above, as well as into 2-day windows equivalent to Fig. 1G. Note that in this case we did not normalize the data since the difference between areas is of interest. Both types of analyses show that V1 null responses do not change relative to preferred responses up to P36. At the same time, PMLS null responses decrease relative to the preferred responses starting around P33. This again strongly argues against small changes in V1 responses that are then amplified in PMLS.

In summary, our data show earlier and faster DS development in PMLS than V1, in a process that involves area-specific decreases in null direction responses as well as increases in preferred responses in both areas.

### Early PMLS development appears specific to DS

Since both V1 and PMLS are tuned for orientation in addition to direction, the development of OS provides the opportunity to assess the areas’ relative development along a second tuning dimension, a crucial test for determining whether PMLS develops early in general, or whether the observations made so far are specific for DSW. OS was quantified using two metrics similar to those for DS, a vector-based metric (Batschelet, 1981; Mazurek et al., 2014) called L_ori_ or 1-CircVar, and an orientation selectivity index (OSI). Again, analyses using either metric showed the same results and only L_ori_ data will be shown here (for OSI results, see Fig. S2 and Table 13).

Intriguingly, the relative development of OS in PMLS versus V1 followed a different pattern than that of DS. In V1, consistent with previous reports (Chapman and Stryker, 1993; Chapman et al., 1996), L_ori_ values were low up to P30. Starting with P31, much higher L_ori_ values could be observed, even if there was significant variability across animals (Fig. 2A). Around P35, V1 L_ori_ values reached the same level as in the young adult group (P40-52: mean L_ori_ 0.53 ± 0.07, median 0.51), which is similar to previous studies in young adults and even in older animals reporting median V1 L_ori_ values of 0.56 (Lempel and Nielsen, 2019), ∼0.5 (Clemens et al., 2012), and ∼0.7 (Popovic et al., 2018). As a consequence of the rapid rise in L_ori_ right around eye opening, dividing the data into the same 3 age groups used for L_dir_ before (P29-32, 33-36, 37+; Fig. 2C) revealed significant differences between the P29-32 group and both other groups (Kruskall-Wallis test and both post-hoc rank-sum tests: p<.01; Table 4), but not between the two older groups.

**Figure 2:**
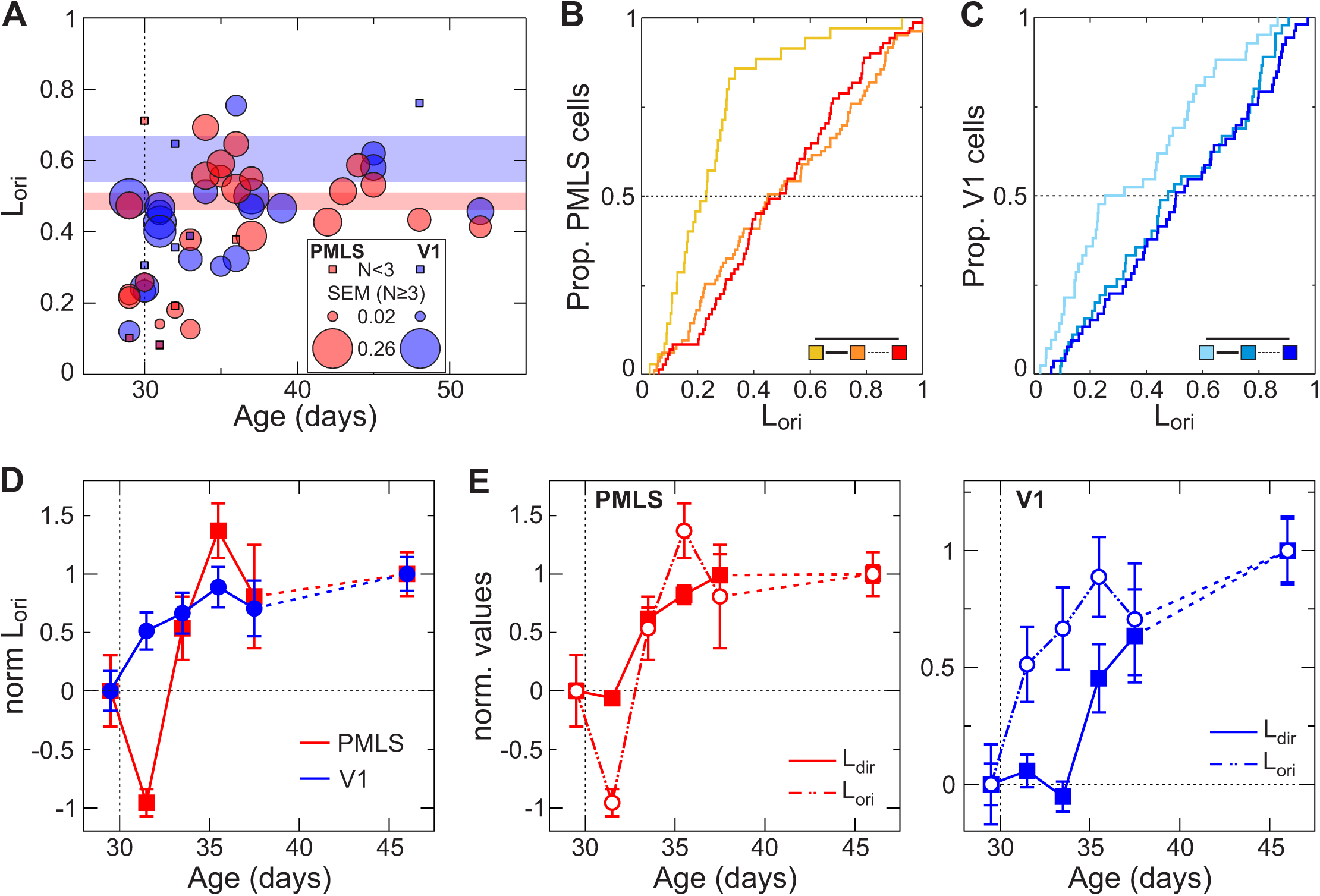
Development of OS in PMLS and V1. (A) Development of L_ori_ on a per-animal basis for V1 and PMLS. As in Fig. 1D, each circle indicates the mean L_ori_ value across all SU that could be isolated in an animal, as a function of the animal’s age. Circle size indicates the SEM for cases with 3 or more neurons. The SEM scale again differs from the scale used for the y-axis and is shown in the legend. Colored bands correspond to mean +/- 1 SEM for animals (P40 and older). (B, C) Cumulative L_ori_ distributions for PMLS and V1. Following the same format as Fig. 1E & F, data were divided into 3 age groups. The legend in the right corner indicates the outcome of rank-sum tests between the different groups (dashed line: p>.01; solid line: p<.01). (D) Normalized L_ori_ as a function of age for both areas. Data were divided into 2-day age bins from P29-38, and an adult bin from P40-52. Bins were normalized (individually for each area) to range from 0 for eye opening to 1 in adult animals. Data for each bin is plotted at the middle of each age bin. Error bars: SEM; same format as Fig. 1G. (E) Normalized L_dir_ and L_ori_ curves for PMLS (left) and V1 (right), replotting the same curves from Fig. 1G and 2D to allow for better comparison of intra-areal development.

**Table 4:** Statistics for Figs. 2B & C.

| Metric | Group | Test | PMLS p value | PMLS other | V1 p value | V1 other |
| --- | --- | --- | --- | --- | --- | --- |
| L<sub>ori</sub> | All age groups | Kruskall-Wallis | <0.001 | X(2)=26.0 | 0.004 | X(2)=10.9 |
| L<sub>ori</sub> | P29-32 vs P33-36 | Rank sum | <0.001 |  | 0.009 |  |
| L<sub>ori</sub> | P29-32 vs P37-52 | Rank sum | <0.001 |  | 0.002 |  |
| L<sub>ori</sub> | P33-36 vs P37-52 | Rank sum | 0.98 |  | 0.6 |  |

In PMLS, OS levels were as low as those found in V1 around P30 (Fig. 2A), with the exception of 2 outliers (one animal contributing 2 neurons, the other 4). Since in all other respects (e.g., their L_dir_ values) these animals were not outliers, their data was included in the OS data set as well. Other than the 2 outliers, OS development in PMLS was delayed relative to V1, with consistent improvements in PMLS L_ori_ values starting only at P34, later than the increase in V1 L_ori_ values. At P34, however, PMLS L_ori_ values immediately assumed the same levels as in the young adult group in a very rapid development. After this point, OS levels appear stable, as the values for the young adult group here (mean L_ori_ 0.5 ± -0.06, median 0.49) are within the range we have previously reported for older animals (mean 0.45 ± 0.04, median 0.45; Lempel and Nielsen, 2019). OS maturation was therefore complete at the same time point in both V1 and PMLS. Indeed, dividing the PMLS data set into the same 3 age groups as above (Fig. 2B) revealed differences only between the P29-32 age groups and both other age groups, comparable to the results for V1 (Kruskall-Wallis test and both post-hoc rank-sum tests: p<.001; see also Table 4).

Given the rapid OS development especially in PMLS, we again turned to binning L_ori_ data into 2-day age windows, normalizing the data to assume 0 and 1 for eye opening and mature levels, respectively, as before (Fig. 2D). Note that the 2 animals with high L_ori_ values around P30 caused the normalized L_ori_ time course for PMLS to initially decrease at P31-32 before rapidly increasing in the next age group. A two-factor ANOVA with age and area (excluding the endpoints used for normalization) was again used to compare development across the areas. It resulted in a significant main effect of age (p<.001; see Table 5), consistent with the increase in OS with age. At same time, there was a significant interaction between area and age (p=.001), due to significantly higher normalized L_ori_ values in V1 in the P31-32 group (post-hoc rank-sum test, p<.001). To rule out possible effects of including the two PMLS outliers in the normalization, we confirmed that a direct comparison of the raw (non-normalized) L_ori_ values for animals aged P31-32 similarly resulted in a significant difference between V1 and PMLS (rank-sum test, p=.003).

**Table 5:**
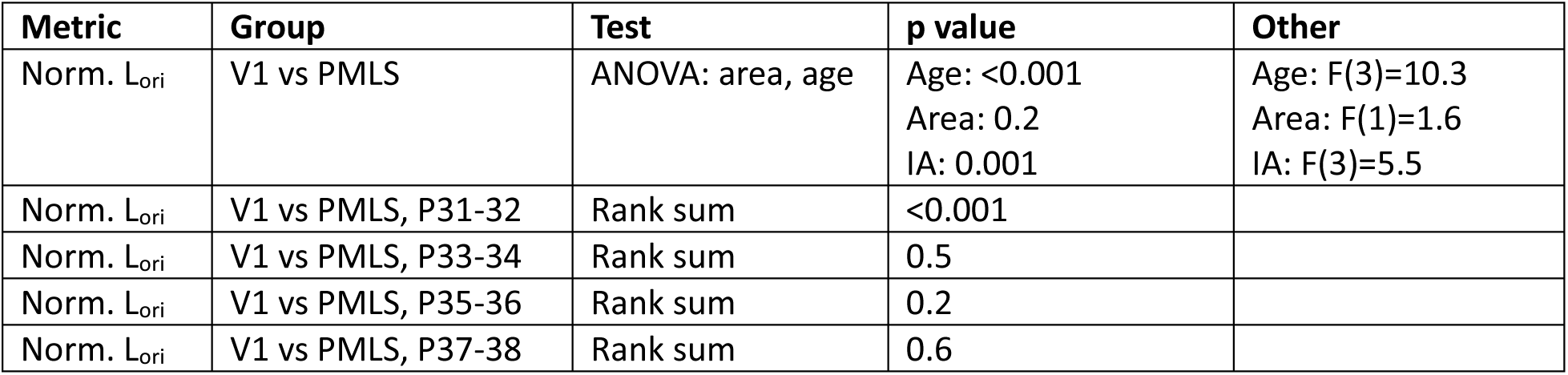
Statistics for Figs. 2D (IA: Interaction)

| Metric | Group | Test | p value | Other |
| --- | --- | --- | --- | --- |
| Norm. L<sub>ori</sub> | V1 vs PMLS | ANOVA: area, age | Age: <0.001<br>Area: 0.2<br>IA: 0.001 | Age: F(3)=10.3<br>Area: F(1)=1.6<br>IA: F(3)=5.5 |
| Norm. L<sub>ori</sub> | V1 vs PMLS, P31-32 | Rank sum | <0.001 |  |
| Norm. L<sub>ori</sub> | V1 vs PMLS, P33-34 | Rank sum | 0.5 |  |
| Norm. L<sub>ori</sub> | V1 vs PMLS, P35-36 | Rank sum | 0.2 |  |
| Norm. L<sub>ori</sub> | V1 vs PMLS, P37-38 | Rank sum | 0.6 |  |

Finally, we also computed changepoints for the non-normalized (but binned) L_ori_ trajectories for each area. In V1, this analysis indicated a change around P31 (95% CI: P31-35; the larger interval reflects the fact that determining the changepoint for V1 L_ori_ is noisier because V1 L_ori_ levels are rising much more gradually with development). The changepoint for PMLS L_ori_ instead occurred at P33 (CI: P33-35), i.e. with a delay relative to V1. Thus, the onset of OS maturation in PMLS lags behind V1, which in turn highlights that the earlier development of PMLS DS is not due to earlier development of the area in general, but that it reflects processes specific to processing of motion direction.

The relative time courses of DS and OS maturation reveal another very intriguing difference between V1 and PMLS development. Comparing these time courses within each area rather than across areas (Fig. 2E), first replicated previous findings of OS development before DS development in V1 (Li et al., 2006). In PMLS, however, our data showed no (or only minimal) delay between DS and OS development (e.g., both normalized time courses show immature levels at P31-32 and significant improvement by P33-34). The normalized L_dir_ and L_ori_ values can again be used to compare the relative development of OS and DS without contamination by overall differences in tuning levels. For V1, a two-way ANOVA with factors age and tuning property on these normalized data resulted in a significant main effect for tuning property (p<.001; see Table 6) and a trend for age (p=.01), but no significant interaction between the factors (p=.4), consistent with differences between the two tuning properties that are maintained throughout development (again, the endpoints used for normalization were excluded from the ANOVA). In contrast, in PMLS only the effect of age reached significance (p<.001), reflecting that DS and OS developed in parallel. The changepoint analysis supports this conclusion as well. As stated above, changepoints for V1 L_ori_ and L_dir_ were P31 and P35, while they fell at P33 for both tuning properties in PMLS. These differences in the sequence of DS and OS development between V1 and PMLS suggest that there might be fundamental differences in the organization of circuits computing direction information in V1 and PMLS (see Discussion).

**Table 6:** Statistics for Fig. 2E (IA: Interaction)

| Area | Group | Test | p value | Other |
| --- | --- | --- | --- | --- |
| V1 | Norm. $L_{ori}$ vs Norm. $L_{dir}$ | ANOVA: age, tuning metric | Age: 0.01<br>Tuning: <0.001<br>IA: 0.4 | Age: F(3)=3.87<br>Tuning: F(1)=12.94<br>IA: F(3)=0.99 |
| PMLS | Norm. $L_{ori}$ vs Norm. $L_{dir}$ | ANOVA: age, tuning metric | Age: <0.001<br>Tuning: 0.4<br>IA: 0.01 | Age: F(3)=17.75<br>Tuning: F(1)=0.73<br>IA: F(3)=3.76 |

### Developmental changes in response latency

In the marmoset, MT has been shown to develop as early as V1, at least anatomically (Bourne and Rosa, 2006). This early development involves a transient, direct pathway from the thalamic pulvinar to MT (Warner et al., 2012; Mundinano et al., 2018). This raises the interesting possibility that early DS development in PMLS is supported by input sources different from those found in adults. Resolving this question will ultimately require anatomical and causal studies. However, response latencies should at least provide some initial insights into network configurations at different ages. As a next step, we therefore decided to analyze the evolution of response latencies in V1 and PMLS with age. Latencies can be quite variable from neuron to neuron. To provide the most robust estimates even at the youngest ages, we increased our sample size by analyzing MU instead of SU data. Here, MU activity is computed as the thresholded but not spike-sorted activity for each recording channel, ensuring that each spike was only counted on a single channel (see Methods). To be included in the latency data set, MU sites had to meet the same visual responsiveness criteria used for SU above (see Table 1 for the resulting data set size). Additionally, we limited the data set to recording sessions in which data was collected simultaneously for V1 and PMLS so that within-animal comparisons were feasible in all cases. This further minimizes confounds in the inter-areal comparisons that could arise from changing response dynamics due to fluctuations in anesthetic state.

For each included MU site, we then determined the onset latency for the site’s best stimulus (as the condition evoking the most robust responses). Latencies were computed by binning spikes into 10 ms bins and computing the first bin in which responses exceeded the mean of pre-stimulus baseline activity by two standard deviations, followed by at least two successive bins exceeding baseline activity (Raiguel et al., 1999). Fig. 3A shows raster plots for 3 example V1 MU sites at different ages, as well as 3 example PMLS sites recorded in the same animals. These examples capture two important trends present in the overall data set shown in Fig. 3B: First, latencies in both areas became significantly shorter with age, until very short latencies were reached in adults (median V1 latency across all MU sites: 60ms; median PMLS latency: 70ms). Second, there was a large delay in PMLS responses relative to V1 in young animals. As latencies in both areas gradually improved over the next days, the gap in latencies between the areas ultimately narrowed and became quite small in adults.

**Figure 3:**
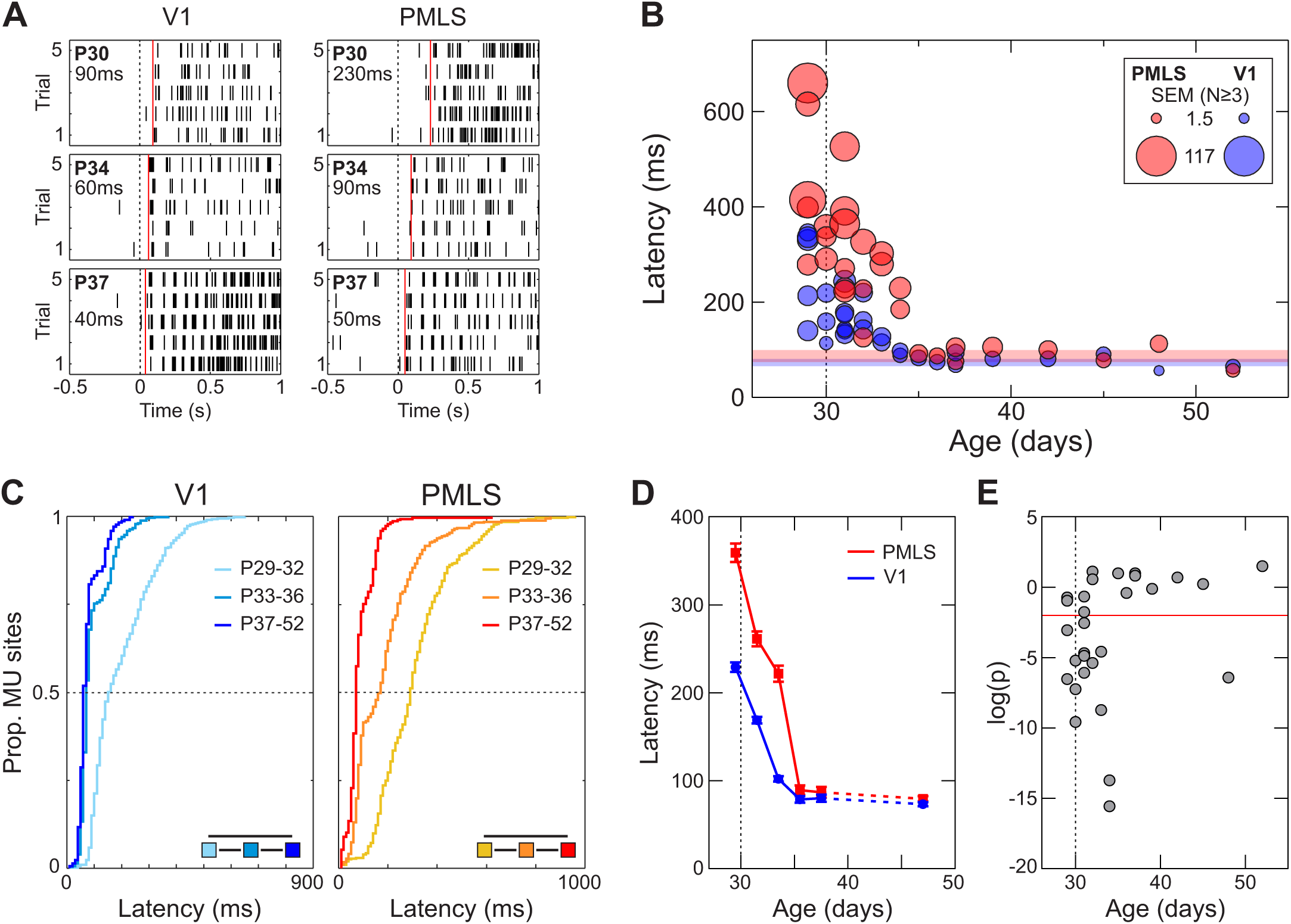
Development of response latency in PMLS and V1. (A) Raster plots for example V1 (left) or PMLS (right) MU sites, recorded at different ages (indicated in upper left in each plot). The dashed line indicates stimulus onset at 0 ms, the red line the latency computed for every site (also listed in each plot). (B) Average MU latency computed for each animal, as a function of the animal’s age. Circle size indicates the SEM for each animal, with the scale for the SEM indicated in the legend. Red and blue band indicate average latency in adult animals (40 days and older), +/- 1 SEM. (C) Cumulative distribution of MU latencies for V1 and PMLS, divided into the standard 3 age groups. The legend summarizes the outcomes of rank-sum tests between groups (solid line: p<.01). (D) Average latency in 2-day age bins from P29-38, and from P40-52, for V1 and PMLS (no normalization). Data points are plotted at the middle of each age bin. Error bars: SEM. (E) p-values resulting from rank-sum test comparing V1 and PMLS latencies for each animal individually. p-values are plotted as a function of the animal’s age, and are Bonferroni-corrected to account for multiple comparisons. The red line indicates a p-value of 0.01.

Fig. 3C & D plot how these findings play out at the per-neuron rather than per-animal level (see Table 7 and Table 8 for full statistics). First, Fig. 3C plots the distribution of latencies for the standard 3 age groups, demonstrating the gradual improvement in latencies with age in both areas (Kruskall-Wallis test across groups per area and all post-hoc rank-sum tests : p<.001). Second, Fig. 3D shows how average latencies, computed in 2-day windows (this time without normalization), evolve with age, highlighting the gradual improvement in latencies between P30 and 35 in both areas, as well as the larger delay in PMLS responses in this earlier phase. Indeed, significant differences between V1 and PMLS emerged only for the three age windows falling between P29 and P34 (2-way ANOVA with factors age and area: all main effects and interaction p<.001; post-hoc rank-sum tests between areas for P29-30, P31-32, P33-34: p<.001), but note that there is still a trend towards larger PMLS latencies for older animals (p<=.05 for P35-36 & animals aged P42-50). Finally, we made use of the fact that the data set was limited to cases with simultaneous recordings in V1 and PMLS to test for each animal individually whether V1 and PMLS latencies differed systematically. For this analysis, the PMLS and V1 latency distributions recorded in the same animal were compared against each other using a rank-sum test, and the resulting p-values were Bonferroni corrected by the total number of animals to account for multiple comparisons. Fig. 3E plots the corrected p-values as a function of each animal’s age, with the red line indicating p=.01. With the exception of one older animal, significant differences between V1 and PMLS latencies using this criterion occurred only before P35.

**Table 7:** Statistics for Fig. 3C.

| Metric | Group | Test |  | PMLS p value | PMLS other | V1 p value | V1 other |
| --- | --- | --- | --- | --- | --- | --- | --- |
| Latency | All age groups | Kruskall-Wallis |  | <0.001 | X(2)=468.3 | <0.001 | X(2)=816.4 |
| Latency | P29-32 vs P33-36 | Rank sum |  | <0.001 |  | <0.001 |  |
| Latency | P29-32 vs P37-52 | Rank sum |  | <0.001 |  | <0.001 |  |
| Latency | P33-36 vs P37-52 | Rank sum |  | <0.001 |  | <0.001 |  |

**Table 8:** Statistics for Fig. 3D (IA: interaction)

| Group | Test | p value | Other |
| --- | --- | --- | --- |
| All age groups | ANOVA: area, age | Area: <0.001<br>Age: <0.001<br>IA: <0.001 | Area: F(1)=193.9<br>Age: F(5)=332.2<br>IA: F(5)=33.6 |
| V1 vs PMLS, P29-30 | Rank sum | <0.001 |  |
| V1 vs PMLS, P31-32 | Rank sum | <0.001 |  |
| V1 vs PMLS, P33-34 | Rank sum | <0.001 |  |
| V1 vs PMLS, P35-36 | Rank sum | 0.05 |  |
| V1 vs PMLS, P37-38 | Rank sum | 0.3 |  |
| V1 vs PMLS, P40-52 | Rank sum | 0.02 |  |

Taken together, our data show that in addition to age-dependent decreases in response latencies in both areas, extra delays occur in PMLS in young animals, possibly reflecting changing input sources and computational demands with age (see Discussion). Interestingly, for PMLS the maturation of latency coincides with the maturation of DS and OS, while for V1 OS develops before latency matures, and DS thereafter.

### Visual experience accelerates DS development in both areas

An important aspect of the development of any visual function is the degree to which it reflects endogenous mechanisms versus the impact of visual experience. In this regard, previous experiments in the ferret have established a highly useful ‘training’ paradigm that allows controlled tests of the impact of different types of visual experience (Li et al., 2008). In this paradigm, visually naïve animals are exposed to a stimulus of interest for a prolonged period of time (while continuously under anesthesia). Baseline tuning levels like DS are determined before and after stimulus exposure to assess whether the chosen training stimulus exerted a measurable influence over neural response properties. For ferret V1, previous studies (Li et al., 2008) reported DS increases after exposure to a drifting grating only – exposure to a gray screen was found to be ineffective, as was exposure to a stationary, flashing grating (a grating with temporal modulation of its contrast). In identifying motion signals as an important cue for V1 DS development, these results demonstrate that the paradigm is a sensitive probe for which types of visual information might influence an area’s development. For this reason, we used it here to gain further insights into PMLS DS development.

Experiments using the training paradigm followed the procedures just described. More precisely, we first determined pre-training tuning levels in V1 and PMLS in visually naïve animals. As before, we used drifting gratings to quantify DS and OS levels. We then exposed the animals to a specific stimulus for 8 h. To minimize adaptation, the paradigm was divided into 20 min long training (or stimulation) blocks in which the stimulus was presented, alternating with 10 min of a blank screen (Fig. 4A). Furthermore, each training block itself consisted of alternating stimulus presentations (5 s) and blank screen periods (10 s). At the conclusion of this training phase, DS and OS levels were again measured in V1 and PMLS and compared to the pre-training levels. Because of the duration of the experiments, we did not attempt to track SU across pre- and post-training measurements. Instead, we assessed DS at the MU level in each case as the more stable measure, reasoning that as electrodes were not moved throughout the experiment, MU activity reflects activity in the same population of neurons before and after exposure (see Methods for inclusion criteria for the training data sets).

**Figure 4:**
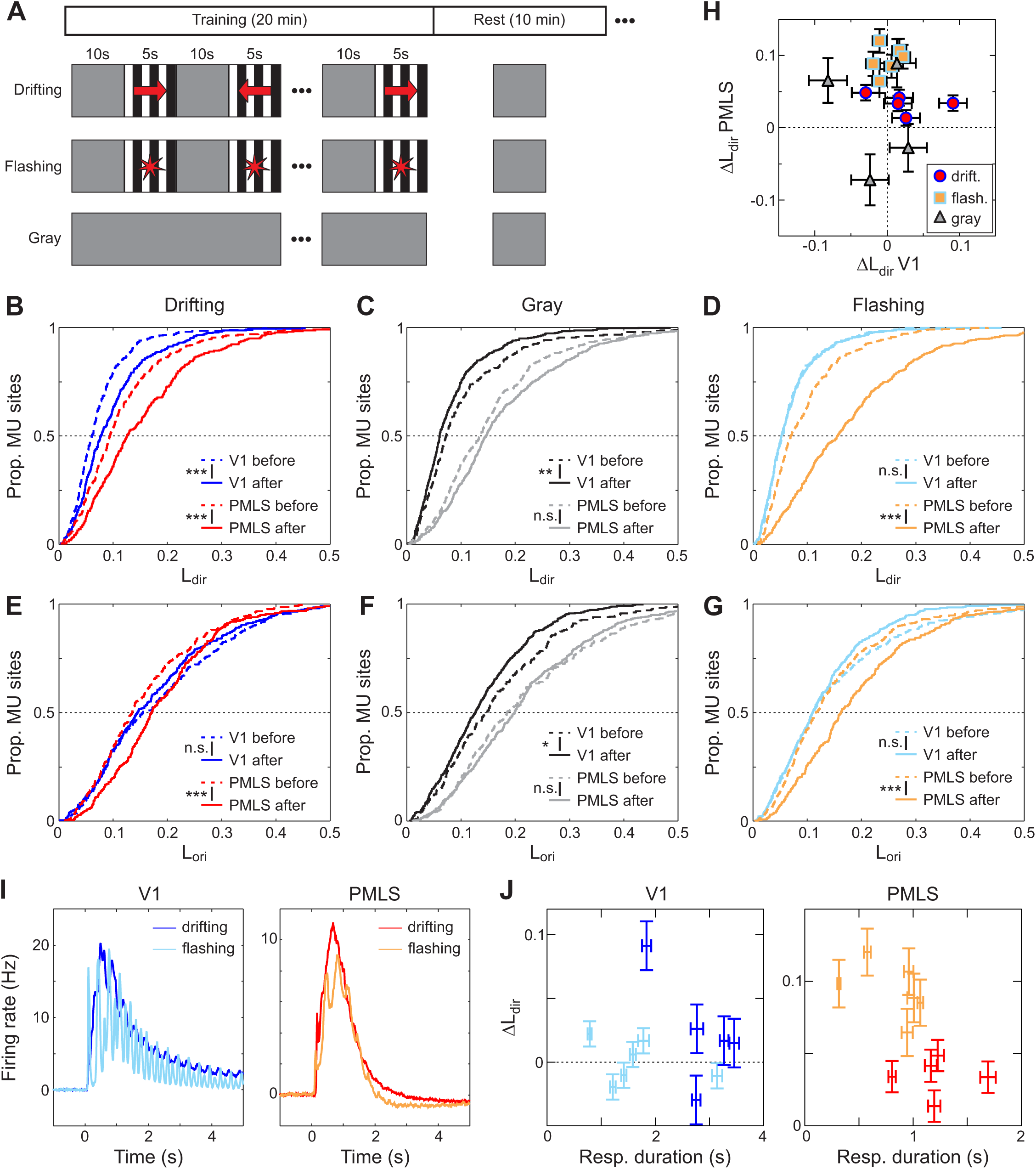
Changes in DS levels in response to different training stimuli. (A) Temporal structure of the training experiments. For two of the 3 cohorts, 20 min-long stimulation blocks were alternated with 10 min-long rest periods for a total duration of 8 h. The stimulation blocks contained alternating sequences of stimulus presentations (5 s) and the gray screen (10 s), and the gray screen was shown during the rest period. In the cohort exposed to drifting gratings, gratings could move in one of two (opposing) directions during the stimulation period. In the cohort exposed to flashing gratings, grating contrast was temporally modulated during each presentation. The third cohort, finally, was only exposed to the gray screen for 8 h. (B-D) Cumulative L_dir_ distributions for V1 and PMLS for MU sites recorded before and after 8 h of exposure to drifting gratings (B), the gray screen (C) or flashing gratings (D). The legends indicate the results of rank-sum tests between pre- and post-training data (**: p<.005, ***: p<.001, n.s.: not significant). (E-G) Cumulative L_ori_ distributions for V1 and PMLS for MU sites, based on the same recordings shown in (B-D), using the same format as (B-D). (H) Correlation in training effect size for L_dir_ between V1 and PMLS. Each symbol corresponds to data from a single animal for which both V1 and PMLS data was available. Effect size is computed for each animal based on the results of the LME models. Colors and symbol shape indicate the stimulus used during the training protocol. Error bars: SE for each per-animal random slope, as estimated by lmefit. (I) Average MU activity evoked by drifting and flashing gratings during the stimulation blocks for V1 and PMLS. Firing rates for each MU site were binned into 10 ms bins and normalized by the average firing rate in the 2 s preceding stimulus onset. Data were then averaged across all stimulus presentations in all 16 blocks, as well as all MU sites and animals. (J) Training effect size as a function of the response duration evoked during the stimulation blocks. Effect size was quantified per animal as in (H). Response duration is computed as the average response duration of all MU sites across all training blocks in an animal. Error bars for the training effect size: SE for each per-animal random slope, as estimated by lmefit. Error bars for the response duration: SEM. Color code as in (H).

As a first test, we investigated the effect of motion cues on tuning development in both areas by using drifting gratings as training stimuli. In each presentation of the training stimulus, the grating could drift in one of 2 opposing directions, so that the axis of motion and orientation was constant throughout. Using this stimulus set, our data replicated the previous findings of increased V1 DS levels after exposure. Fig. 4B plots the cumulative L_dir_ distribution across MU sites before and after training, demonstrating a significant increase in V1 DS levels after training (rank-sum test, p<.001). While this increase is consistent with previous studies, the effect size appears to be significantly smaller in our animals than in those earlier studies (Li et al., 2008; Van Hooser et al., 2012; Ritter et al., 2017; Roy et al., 2020). A possible explanation for this discrepancy is a difference in the amount of natural visual experience per animal across studies. None of the animals studied here had naturally opened their eyes at the start of the training experiments. In contrast, previous studies included animals of the same age range, but with 1-3 days of visual experience (Van Hooser et al., 2012; van Hooser, personal communication). Indeed, the mean pre-training V1 L_dir_ level was higher in at least one of these previous studies than in our sample, as would be expected in animals with some visual experience (here: mean pre-training V1 L_dir_ across all training cohorts: 0.07 +/- 0.001 (SEM); Ritter et al., 2017: 0.105 +/- 0.017). It is plausible that even brief amounts of natural visual experience can predispose V1 circuits to be able to change more rapidly in response to the training stimuli, therefore resulting in larger effects. Also note that previous studies using the same training paradigm have shown that changes in DS are strongest at the trained direction in V1, both because neurons initially preferring this direction strengthen in their responses, but also because stimulus preferences become aligned to the training stimulus (Li et al., 2008; Van Hooser et al., 2012). Here, we chose the training direction to drive responses at both recording sites, but did not aim to match them precisely to stimulus preferences at either location. It is possible that some variability in the training effect size is caused by a different degree of alignment between the training direction and the stimulus preferences at our recording locations.

At the same time, since the effect size in our experiments is smaller, there might be a concern that the population data mostly reflects changes in a single animal. In addition to analyzing data across animals, we therefore also used a linear mixed-effect (LME) model to directly control for grouping of data by animal. More precisely, we fit the before- and after-training L_dir_ data with an LME model with a fixed effect for training phase (before/after training), allowing random intercepts and slopes per animal (see Methods and Fig. S3). On the full data set, the effect of training phase indeed was not significant (p=.2). However, 4 out of the 5 animals showed an increase in L_dir_ values after training (Table 9). If the 5^th^ animal (which showed a decrease) is excluded, the effect of training phase becomes significant (p=.009).

**Table 9:** LME results for Fig. 4B-D (Phase: Slope for the fixed effect training phase (before/after); Slope: fixed effect slope + random effect slope per animal)

| Training stim. | Parameter | V1 | PMLS |
| --- | --- | --- | --- |
| Drifting, $L_{dir}$ | Phase: Estimate | Full data: 0.02<br>Reduced data: 0.04 | 0.03 |
| Drifting, $L_{dir}$ | Phase: p value | Full data: 0.2<br>Reduced data: 0.009 | <0.001 |
| Drifting, $L_{dir}$ | Slope per animal | 0.02, 0.09, -0.03, 0.02, 0.03 | 0.04, 0.03, 0.05, 0.03, 0.01 |
| Gray, $L_{dir}$ | Phase: Estimate | -0.03 | 0.01 |
| Gray, $L_{dir}$ | Phase: p value | 0.2 | 0.7 |
| Gray, $L_{dir}$ | Slope per animal | -0.10, 0.01, -0.02, 0.03, -0.08 | 0.09, -0.07, -0.002, -0.03, 0.07 |
| Flashing, $L_{dir}$ | Phase: Estimate | 0.0008 | 0.09 |
| Flashing, $L_{dir}$ | Phase: p value | 0.9 | <0.001 |
| Flashing, $L_{dir}$ | Slope per animal | -0.02, 0.02, 0.02, -0.01, -0.01, 0.006 | 0.09, 0.11, 0.10, 0.08, 0.12, 0.06, 0.09 |

Importantly, our data show for the first time that exposure to drifting gratings also has a measurable impact on PMLS DS levels. Combined across all animals, L_dir_ levels significantly increased in PMLS with training (rank-sum test, p<.001). The LME model similarly returned a significant effect of training phase (p<.001), and increases in L_dir_ values for all 5 animals (Table 9). Thus, it appears that drifting gratings contain the necessary information to drive DS development not just in V1, but also in PMLS.

To support this conclusion, it is important to compare the results to the effects of exposing animals to a blank screen for 8 h, to determine whether the observed DS changes indeed were due to information contained in the stimulus versus a more general effect of unspecific visual stimulation (since the eyes are opened at the start of the experiment) or simply the passage of time. Fig. 4C and S3 plot the L_dir_ changes observed before and after exposure to a homogenously gray screen for 8 h for both V1 and PMLS. Based on analyzing the population data as a whole, V1 L_dir_ levels actually decreased during this time period (rank-sum test, p=.003). In PMLS, a trend towards higher L_dir_ levels did not reach significance (rank-sum test, p=.01). The LME models, which account for per-animal effects, showed no significant change in the L_dir_ values for either area (V1: p=.2, PMLS: p=.7), and positive and negative L_dir_ changes occurred with the same frequencies across animals for both areas (Table 9). Thus, for PMLS as for V1, the DS changes induced by the drifting grating indeed reflected the impact of the particular visual information presented during the training, not unspecific effects.

In addition to changes in DS, we also tested whether OS levels were affected by the training paradigm (Fig. 4E-F & S3). In V1, increases in OS were previously observed after exposure to a drifting grating, but also after exposure to a gray screen (Li et al., 2008; Ritter et al., 2017), suggesting that these improvements occur independent of the particular visual information provided. Here, we observed no changes in V1 L_ori_ levels after exposure to a drifting grating across the full data set (rank-sum test, p=.3; LME model: p=.8, Table 10). However, the same animal that was an outlier in terms of L_dir_ changes in this cohort also was the only animal that showed decreases in L_ori_ after training (see LME slopes in Table 10). Excluding this animal resulted in significant changes in V1 L_ori_ levels before and after training (population level rank-sum test, p<.001; LME model: p=.002, Table 10). In contrast, exposure to a gray screen actually resulted in a small but significant decrease in V1 L_ori_ at least at the per-site level in our experiments (rank-sum test across all MU sites, p=.006; LME model: p=.08; Table 10). Thus, in our data set the V1 OS changes are specific to the stimulus used, not unspecific as shown previously. Again, the differences between our and previous results may lie in the difference in starting point. In this context it is worth noting that, just like baseline DS levels, baseline OS levels were lower in our sample than in previous studies (here: mean pre-training V1 L_ori_ across all cohorts: 0.18 +/- 0.002; Ritter et al: 0.383 +/-0.036).

**Table 10:** LME results for Fig. 4E-G (Phase: Slope for the fixed effect training phase (before/after); Slope: fixed effect slope + random effect slope per animal)

| Training stim. | Parameter | V1 | PMLS |
| --- | --- | --- | --- |
| Drifting, $L_{ori}$ | Phase: Estimate | Full data: -0.009<br>Reduced data: 0.03 | 0.03 |
| Drifting, $L_{ori}$ | Phase: p value | Full data: 0.8<br>Reduced data: 0.002 | <0.001 |
| Drifting, $L_{ori}$ | Slope per animal | 0.06, 0.01, -0.17, 0.04, 0.03 | 0.02, 0.04, 0.02, 0.04, 0.03 |
| Gray, $L_{ori}$ | Phase: Estimate | -0.05 | -0.05 |
| Gray, $L_{ori}$ | Phase: p value | 0.08 | 0.4 |
| Gray, $L_{ori}$ | Slope per animal | -0.15, 0.002, -0.07, 0.02, -0.05 | -0.01, -0.24, 0.05, -0.12, 0.08 |
| Flashing, $L_{ori}$ | Phase: Estimate | -0.02 | 0.05 |
| Flashing, $L_{ori}$ | Phase: p value | 0.3 | 0.005 |
| Flashing, $L_{ori}$ | Slope per animal | 0.02, -0.009, 0.03, -0.03, -0.03, -0.13 | -0.009, 0.03, 0.09, 0.02, 0.06, 0.06, 0.09 |

In PMLS, L_ori_ levels improved significantly after exposure to drifting gratings (rank-sum test, p<.001; LME model: p<.001; 5/5 animals with positive slope; Table 10). No change was observed in the control cohort exposed to a gray screen (rank-sum test, p=.8; LME model: p=.4; Table 10), indicating that in PMLS, changes in OS, like changes in DS, were largely driven by the stimulus information provided during training.

### PMLS DS development is sensitive to more visual cues than V1

The training experiments so far demonstrate that patterned visual experience can impact not just V1 but also PMLS DS (and OS) development. The next question then is whether certain types of visual information are more important in this process than others. As mentioned above, exposure to a training stimulus lacking motion signals previously proved ineffective in changing V1 DS levels. To probe whether motion signals are similarly crucial for PMLS DS development, we performed a third set of training experiments, in which animals were exposed to flashing gratings. These gratings were generated by temporally modulating grating contrast between 0% and 100%. Importantly, no contrast reversals were included. This ensured that while the stimulus contrast varied dynamically, the spatial position of the stimulus remained fixed, and no motion cues were provided.

The results of these experiments are plotted in Fig. 4D and Fig. S3. In agreement with the earlier study by Li et al. (Li et al., 2008), V1 L_dir_ levels remained unchanged after 8 h of exposure to the flashing grating (rank-sum test, p=.9; LME model: p=.9, 3/6 animals with L_dir_ increases; Table 9). In contrast, the same stimulus was highly effective in PMLS, with significant improvements in L_dir_ at the end of the training experiment (rank-sum test, p<.001, LME model: p<.001, 7/7 animals with L_dir_ increases; Table 9). In fact, at least in the cohorts studied here, flashing gratings resulted in larger changes in PMLS L_dir_ levels than drifting gratings. We used the LME models to estimate training effect size per animal (see Methods). Based on this metric, training effects were significantly larger for flashing than drifting gratings in PMLS (rank-sum test, p=.003). Our data therefore strongly suggest that contrary to V1, explicit motion cues were not required for promoting DS development in PMLS. While the flashing gratings are stationary spatially, they nonetheless dynamically changed over time. Based on our data, PMLS is sensitive to these temporal cues, while V1 is not (or to a much lesser degree). It remains to be determined whether temporal cues alone are strong enough that a spatially uniform stimulus that changes luminance in time, but lacks spatial information, would nonetheless be sufficient in causing PMLS DS increases.

In addition to highlighting the role of different types of visual information, the differential impact of flashing gratings on V1 and PMLS represents a striking example of dissociation in DS development between the two areas reminiscent of the observations during normal development. The training data generally provide an opportunity to further test this idea, because they can be used to ask whether L_dir_ changes induced by the training protocol in V1 are correlated with L_dir_ changes in PMLS in the same animal. Fig. 4H plots the L_dir_ changes in V1 against the changes in PMLS for each animal (with training effect sizes determined from the per-animal LME models as above). Note that for this plot, we only used animals in which data from both areas were available. The lack of correlation of effect sizes is evident in this figure (r=-0.09, p=.8), and supports the notion that DS development in PMLS does not track V1 DS development.

Lastly, the difference in response to the flashing gratings between V1 and PMLS also extended to OS (Fig 4G & S3). In PMLS, L_ori_ levels increased after training (rank-sum test, p<.001; LME model: p=.005; 6/7 animals with L_dir_ increases; Table 10), while no change was detected in V1 (rank-sum test, p=.2; LME model: p=.3; Table 10). As for L_dir_, there was no correlation between the changes in V1 and PMLS L_ori_ levels across animals (Fig. S4A; r=0.01, p=.97).

As intriguing as the diverging effects of the flashing gratings on V1 and PMLS are, it needs to be ruled out that the flashing gratings simply fail to activate V1, therefore rendering the stimulus ineffective in driving L_dir_ change. To address this question, we turned to MU responses recorded during the stimulus presentation periods in the training blocks, and compared responses to flashing gratings against those to drifting gratings as an effective training stimulus. Fig. 4I plots the average response time courses evoked by both types of stimuli for the two areas (averaging across all blocks and all MU sites in all animals for the 2 training cohorts). These time courses clearly show that flashing gratings did not fail to drive V1, but rather that both types of stimuli evoked reliable activity in both areas. In fact, while peak rates differed between the two areas (Fig. S5), they did not differ between stimulus types (two-way ANOVA with factors area and stimulus type; area: p<.001, type: p=.1, interaction: p=.3; Table 11). The major difference between the stimulus types appeared to be the duration of the evoked response (quantified as the time with firing rates above baseline; see Methods), and as a consequence the average rates (Fig. S5, Table 11). More precisely, response durations were significantly shorter for flashing than drifting gratings in both areas (ANOVA with factors area and stimulus type, both main effects and interaction p<.001; individual rank-sum tests per area: p<.001; Table 11). Interestingly, the difference in response duration was more pronounced in V1 than PMLS. However, if lower V1 drive due to shorter response durations indeed was the reason for a lack of DS change after training with flashing gratings, then response duration should be correlated with the training-induced change in L_dir_. As is obvious from Fig. 4J, in V1 this is not the case for both stimuli combined or individually (both conditions combined: r=-0.07, p=.8; drifting: r=-0.7, p = .2; flashing: r=-0.3, p = .5). The same is the case for PMLS (both conditions: r=-0.6, p=.04; drifting: r=0.001, p=1; flashing: r=-0.5, p=.3). Thus, the different outcomes of exposure to flashing gratings for V1 and PMLS tuning levels cannot simply be attributed to a failure to drive V1 response, but indeed appear to reflect sensitivity to different stimulus attributes in the two areas.

**Table 11:** Statistics for Fig. S5 (IA: interaction). All rank-sum tests compare drifting versus flashing gratings.

| Area | Metric | Test | p value | Other |
| --- | --- | --- | --- | --- |
| V1 & PSS | Peak rate | ANOVA: area, stimulus type | Area: <0.001<br>Stim type: 0.12<br>IA: 0.3 | Area: F(1)=282.4<br>Stim type: F(1)=2.4<br>IA: F(1)=1.2 |
| V1 & PMLS | Response duration | ANOVA: area, stimulus type | Area: <0.001<br>Stim type: <0.001<br>IA: <0.001 | Area: F(1)=973.9<br>Stim type: F(1)=400.4<br>IA: F(1)=93.7 |
| V1 & PMLS | Average rate | ANOVA: area, stimulus type | Area: <0.001<br>Stim type: <0.001<br>IA: <0.001 | Area: F(1)=312.0<br>Stim type: F(1)=63.7<br>IA: F(1)=18.5 |
| V1 | Response duration | Rank sum | <0.001 |  |
| V1 | Average rate | Rank sum | <0.001 |  |
| PMLS | Response duration | Rank sum | <0.001 |  |
| PMLS | Average rate | Rank sum | <0.001 |  |

As a final check, we compared the baseline levels of selectivity pre-training across all training cohorts for both areas. Animals in the 3 cohorts were matched in terms of age range (drifting gratings: P29-33, mean 30.8; gray screen: P28-31, mean 29.8; flashing gratings: P29-32, mean 30.6), and their lack of visual experience. Nonetheless, it is possible that a different starting point (in terms of tuning development) might predispose some animals for larger training effects than others. We indeed observed systematic differences in the baseline DS and OS levels across the 3 cohorts. At the same time, these differences did not appear to be related to the differences in training effects. At the per-site level, V1 L_dir_ levels were significantly higher in the cohort exposed to a gray screen than both other groups (rank-sum test, p<.001 for both tests; Table 12), and there was a trend toward higher L_dir_ levels in the cohort trained with drifting gratings than that trained with flashing gratings (rank-sum test, p=.03, Table 12). The cohort exposed to drifting gratings therefore showed baseline tuning levels that fell between both other groups. Nonetheless, this was the only group that showed improvements in DS with training, arguing against a simple relationship between baseline tuning levels and training effects. When analyzed at the per-animal level using the pre-training mean and training effect slope determined from the LME (Fig. S4B), a negative correlation between baseline tuning levels and training effect size arose across all 3 groups (r=-0.7, p=0.003). This correlation is driven by the results in the cohort exposed to the gray screen: Excluding the gray screen cohort from the analysis results in a non-significant correlation (r=0.2, p=.5), as is the correlation for each group on its own (drifting: r=0.5, p=.4; flashing: r=0.1, p=.8; gray: r=-0.9, p=.02). The per-animal data therefore also argue that the differences in effectiveness of different training stimuli do not merely reflect different starting conditions across the cohorts.

**Table 12:** Comparisons of baseline L_dir_ and L_ori_ levels across cohorts for V1 and PMLS (related to Fig. S4).

| Metric | Group | Test | PMLS p value | PMLS other | V1 p value | V1 other |
| --- | --- | --- | --- | --- | --- | --- |
| $L_{dir}$ | All cohorts | Kruskall-Wallis | <0.001 | $X(2)=93.0$ | <0.001 | $X(2)=30.1$ |
| $L_{dir}$ | Drifting vs gray | Rank sum | <0.001 | | <0.001 | |
| $L_{dir}$ | Drifting vs flashing | Rank sum | <0.001 | | 0.03 | |
| $L_{dir}$ | Flashing vs gray | Rank sum | <0.001 | | <0.001 | |
| $L_{ori}$ | All cohorts | Kruskall-Wallis | <0.001 | $X(2)=77.1$ | <0.001 | $X(2)=22.5$ |
| $L_{ori}$ | Drifting vs gray | Rank sum | <0.001 | | 0.1 | |
| $L_{ori}$ | Drifting vs flashing | Rank sum | 0.02 | | <0.001 | |
| $L_{ori}$ | Flashing vs gray | Rank sum | <0.001 | | 0.003 | |

**Table 13:**
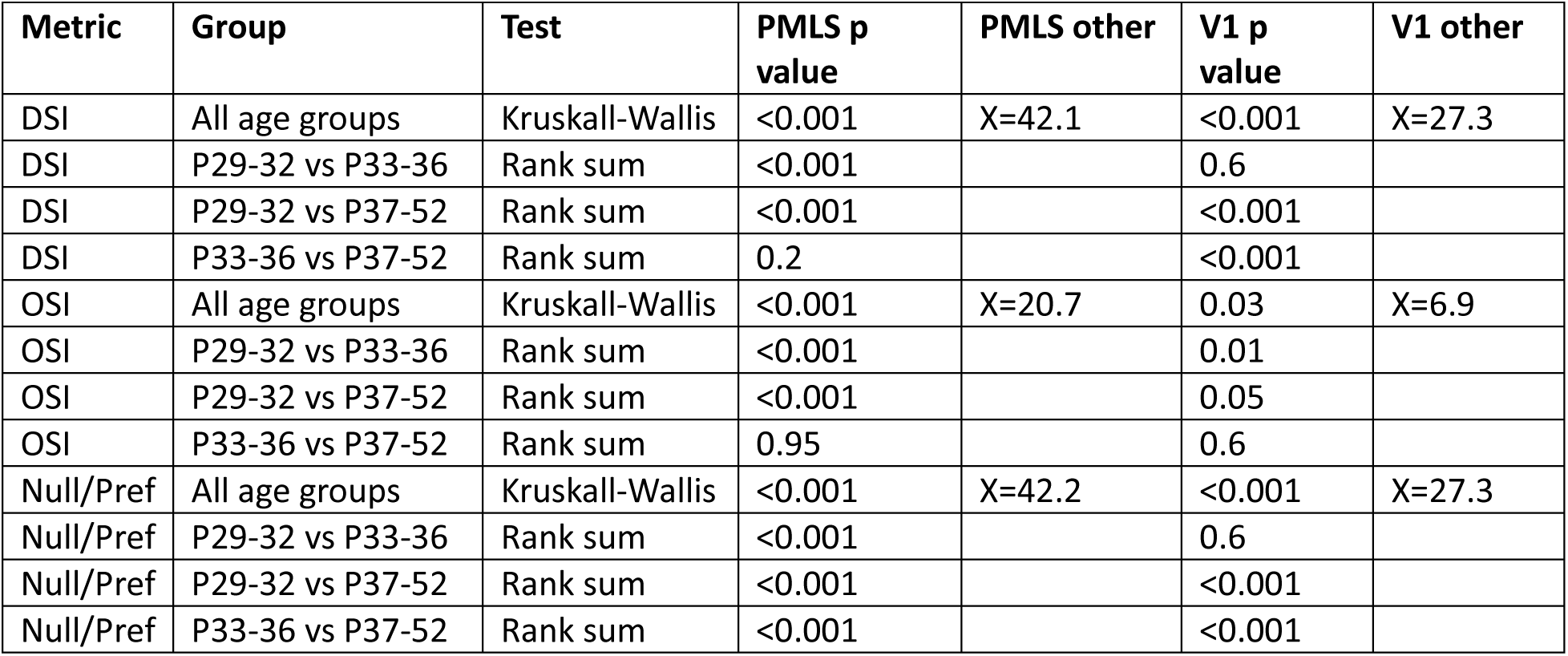
Statistics for Figs. S2B, C, E-H.

| **Metric** | **Group** | **Test** | **PMLS p value** | **PMLS other** | **V1 p value** | **V1 other** |
| --- | --- | --- | --- | --- | --- | --- |
| DSI | All age groups | Kruskall-Wallis | <0.001 | X=42.1 | <0.001 | X=27.3 |
| DSI | P29-32 vs P33-36 | Rank sum | <0.001 |  | 0.6 |  |
| DSI | P29-32 vs P37-52 | Rank sum | <0.001 |  | <0.001 |  |
| DSI | P33-36 vs P37-52 | Rank sum | 0.2 |  | <0.001 |  |
| OSI | All age groups | Kruskall-Wallis | <0.001 | X=20.7 | 0.03 | X=6.9 |
| OSI | P29-32 vs P33-36 | Rank sum | <0.001 |  | 0.01 |  |
| OSI | P29-32 vs P37-52 | Rank sum | <0.001 |  | 0.05 |  |
| OSI | P33-36 vs P37-52 | Rank sum | 0.95 |  | 0.6 |  |
| Null/Pref | All age groups | Kruskall-Wallis | <0.001 | X=42.2 | <0.001 | X=27.3 |
| Null/Pref | P29-32 vs P33-36 | Rank sum | <0.001 |  | 0.6 |  |
| Null/Pref | P29-32 vs P37-52 | Rank sum | <0.001 |  | <0.001 |  |
| Null/Pref | P33-36 vs P37-52 | Rank sum | <0.001 |  | <0.001 |  |

In PMLS, we similarly observed a systematic difference in baseline L_dir_ levels across the 3 cohorts (Kruskall-Wallis test, p<.001; Table 12). Here, tuning levels were lowest in the cohort exposed to flashing gratings when analyzed at the population level, followed by the drifting grating cohort and the gray screen cohort (rank-sum tests, all p<.001; Table 12). When analyzed at the per-animal level, we consequently observed a trend for a negative correlation between pre-training means and the training effect slope across all 3 cohorts (r=-0.4, p=.08; Fig. S4D). However, as for V1 this correlation was driven by the gray screen cohort, and correlations were not significant once that cohort was excluded from the analysis (r=-0.2, p=.6) and for each cohort individually (drifting: r=0.8, p=.1; flashing: r=0.1, p=.8; gray: r=-0.2, p=.7). Thus, again, it seems highly unlikely that the effectiveness of different stimuli on PMLS tuning purely reflects different baseline tuning levels. L_ori_ results, finally, generally paralleled the results for L_dir_ for both areas, and similarly point to stimulus content, rather than baseline tuning levels, as the major source for different training outcomes (Fig. S4C & E, Table 12).

The following reasons further support the notion that the consistent negative correlations we observed between baseline levels and training effects are spurious rather than a real effect: First, a negative correlation between baseline level and training effect size might be expected if baseline levels were close enough to mature levels to limit possible training effect sizes (i.e., causing a ceiling effect). In all cases here, baseline levels were immature enough that even after training, tuning levels were below the mature values, making it unlikely that a ceiling effect caused the negative correlation. Second, it is possible that higher baseline levels indicate a more mature developmental state that is less susceptible to the training paradigm. As discussed above, a comparison of our results with those of previous studies shows that to the contrary, larger effects are observed in animals with more visual experience and higher baseline tuning levels that found in our data, arguing against this explanation (see also Discussion).

## Discussion

A key feature of the experiments presented here is the direct comparison of multiple stages of the ferret’s motion pathway. This network-level perspective reveals a complex picture of development, one that does not agree with a simple feedforward sweep in which early stages develop before later ones. Neither are our data consistent with an equally simple reversal of the hierarchy early in life, in which PMLS becomes the primary driver. DS develops earlier in PMLS than in V1, in violation of a strictly feedforward sequence. Yet, PMLS OS does not develop early, and neither does response latency. Paired with different sensitivity to visual cues, it is clear that the developmental status of one area cannot be predicted by that of the other, This highlights the necessity to study the developing visual system not one area at a time, but by probing multiple network nodes simultaneously to adequately capture the status of all interacting areas.

The differences in V1 and PMLS development observed here raise the intriguing possibility that at least some PMLS functions develop independently from V1, possibly supported through inputs bypassing V1. While the comparison of tuning functions across areas presented here suggests this conclusion, it will require causal manipulations to actually test the influence of one area on the development of the other. In general, since our data are restricted to measurements of spiking output from both areas, they cannot identify the mechanisms that support the DS development in either area. As discussed above, our data show stable V1 responses during time periods of rapid PMLS DS development, arguing against a model in which subtle differences in V1 responses are simply enhanced in PMLS, and instead are more consistent with either PMLS-internal mechanisms, and/or input from other sources. Yet, direct measurements of V1 input to PMLS neurons will be needed to confirm this hypothesis.

A possible role of V1-bypassing inputs in PMLS development is especially intriguing because of developmental patterns observed for the primate motion pathway. Specifically, marmoset MT develops structurally as early as V1 (Bourne and Rosa, 2006), consistent with our observation of the early functional development of PMLS. This early MT development is supported by a transient pathway providing retinal input to MT through the thalamic pulvinar, different from the adult flow of information through the lateral geniculate nucleus (LGN) and V1 to MT (Warner et al., 2012; Mundinano et al., 2018). Our data suggest this might also occur in the carnivore motion pathway, possibly because early development of basic motion functions is important for survival (Warner et al., 2012). So far, only the development of circuits linking V1 and PMLS has been investigated in ferrets. These studies suggest that around eye opening, interactions between V1 and PMLS are privileged over the interactions of other cortical areas and V1: In this age group, both feedforward and feedback projections between V1 and PMLS are stronger than the same projections between V1 and neighboring areas 18 and 19, a pattern that reverses later in life (Khalil and Levitt, 2014; Khalil et al., 2018).

However, the importance of the V1-PMLS loop over other cortical loops does not preclude the existence of strong subcortical input to PMLS, possibly undergoing age-dependent changes similar to the observations in primates. While no anatomical data exist at this time that could address this question, our latency data are quite interesting in this regard: Response latencies in both areas shorten with age, something that has also been observed for monkey V1 and V2 and might suggest general circuit maturation (Zhang et al., 2008). At the same time, while there is considerable overlap in V1 and PMLS latencies in adults similar to adult primate V1 and MT (Raiguel et al., 1989, 1999; Schmolesky et al., 1998), up to P35 latencies are significantly slower in PMLS than V1. An intriguing hypothesis then is that the extra delay reflects the cost incurred by driving PMLS activity through different input sources compared to adults (interesting candidates include LGN, pulvinar or superior colliculus). Since DS is sparse in ferret LGN at least from eye opening on (Murphy et al., 2020; Stacy et al., 2023), the extra delay may also reflect that different input sources require PMLS to compute direction signals *de novo*, rather than relying on ‘precomputed’ direction signals provided by V1. While further experiments are needed to answer these questions, the available developmental data from ferrets and primates nonetheless strongly argue that development of the visual hierarchy is best described as a dynamic sequence of unique networks that can be configured quite differently from the adult, rather than a gradual, layer-by-layer emergence of the adult hierarchy (Bourne et al., 2024). Significant reconfiguration of the visual hierarchy with development has also been observed in mice, in that higher areas associated with dorsal versus ventral stream-like processing mature at different rates (Murakami et al., 2017; Smith et al., 2017), and the connectivity between thalamus, V1 and higher visual areas undergoes drastic changes with development (Murakami et al., 2022), pointing to the universality of this observation.

One of the unexpected results of our study is the different impact of visual cues on V1 and PMLS DS development. Motion computations are based on the joint processing of spatial and temporal signals. Our data, as well as the results of previous studies, show that only stimuli containing both cues (i.e., real motion) are efficient in promoting the development of V1 DS circuits. This holds both in animals pre-eye opening (our data), as well as animals a few days post-eye opening (Li et al., 2008). At the same time, our data show that drive arising from temporal transients only is sufficient for DS development in PMLS. In fact, at least in our experiments flashing gratings were more effective than stimuli showing actual motion. In addition to pointing to different mechanisms shaping the developing circuits in the two areas, these different sensitivity profiles also suggest that alterations in visual input during development might cause very different outcomes for V1 and PMLS, as long as they impact spatial and temporal content differently.

As discussed above, the DS (and OS) improvements in V1 are smaller here than reported previously even for an effective stimulus, likely due to differences in visual experience between studies. While this raises the possibility that our results underestimate the impact of different stimulus dimensions on V1 development, it should be noted that flashing stimuli remained ineffective in promoting V1 DS even 1-3 days after eye opening (see above). Importantly, it also does not impact the conclusion that PMLS development differs from that of V1. In fact, the robust effects in PMLS paired with smaller V1 effects seen in our data, compared to larger V1 effects seen after more visual experience previously, imply that stimulus-driven plasticity is effective earlier in PMLS than V1. A similar argument can be made regarding the concern that differential effects of anesthesia might have impacted the training outcomes. Since it is generally assumed that anesthesia affects higher processing stages more than lower ones, smaller effects in PMLS than V1 would be predicted based on anesthesia effects, inconsistent with our findings. To explain our data, one would instead have to assume that PMLS is less affected than V1, effectively suggesting that it occupies a lower rank in the visual hierarchy than V1 around eye opening.

In addition to these insights into visual hierarchy development, our data may also provide valuable clues regarding motion processing in adult cortex. One important observation in this respect are the relative developmental time course of OS and DS. In V1, we and others observe OS development before DS (Li et al., 2006; Clemens et al., 2012). At least in the ferret, this sequence is reflected in the functional organization of the area: Orientation maps develop first, and show minimal discontinuities in the form of pinwheels (Chapman et al., 1996; Weliky et al., 1996; Li et al., 2006). The later developing direction maps are then constrained by the pre-existing orientation maps. As a consequence, they are nestled inside the orientation maps, and the discontinuities for the direction map take the form of linear fractures with direction shits of 180 deg (Weliky et al., 1996). In PMLS, DS development begins before OS development, but both mature at the same time. Not only does this demonstrate that PMLS DS development is not purely a consequence of OS improvements, it raises the interesting possibility that the PMLS functional architecture is optimized for direction rather than orientation.

Another relevant observation for adult PMLS function is the decrease in null responses concurrent with DS development. Null direction inhibition is thought to play a role in the generation of DS in primate MT (Livingstone et al., 2001; Thiele et al., 2004, 2012), and we have suggested the same might be the case in ferret PMLS (Lempel and Nielsen, 2019). While the anatomical and functional development of inhibitory neurons in ferret V1 has received some attention (e.g., Gao et al., 1999, 2000; Mulholland et al., 2021; Chang and Fitzpatrick, 2022), there are currently no comparable data for PMLS. Further studies are therefore necessary to resolve whether the decreased null responses are indeed the consequence of developing inhibition (rather than e.g. a change in feedforward drive). A further open question concerns the link between development of basic DS versus more complex PMLS tuning properties, including the role of different types of visual information. We have previously shown that selectivity for the global motion of a pattern matures in PMLS around P42, i.e. about a week after the maturation of DS. Our previous data also point to a general role of visual experience in this process (Lempel and Nielsen, 2021). Whether different visual cues play different roles however is currently unknown, but might be addressable using the training paradigm, in addition to testing whether pattern selectivity can be induced at the same time as DS or only after. As we did not quantify the degree of pattern motion selectivity in PMLS after the training experiments, these remain questions for future experiments.

Finally, the complex nature of the relative V1-PMLS development we observed will be important to consider in the context of both developmental disorders and adult plasticity mechanisms. Early development, paired with sensitivity to a broad range of visual cues, likely means increased vulnerability to disruptions of normal development early in life – consistent with the observation that motion vision is impaired in a range of developmental disorders (see e.g. Atkinson, 2017; Bourne et al., 2024 for reviews). On the other hand, circuit mechanisms that support early DS development in PMLS may still be available for reactivation later in life and provide mechanisms for phenomena like blindsight (Weiskrantz et al., 1995; Weiskrantz, 1996) or the ability to regain some visual function after V1 stroke through training on a motion task (Huxlin et al., 2009).

## Conflict of interest

The authors declare no competing financial and non-financial interests.

## Acknowledgments

This work was funded by NIH EY027853 (KJN) and NIH EY035807 (KJN). We are thankful for O. Garalde, R. Vistein and other members of the Nielsen lab for experimental support, and thank J. Killebrew, W. Nash and W. Quinlan for their technical support.

## Notes

### Competing Interest Statement

The authors have declared no competing interest.

### Summary of Updates

New data and analyses were added to the manuscript.

## References

1. Atkinson J (2017) The Davida Teller Award Lecture, 2016: Visual Brain Development: A review of “Dorsal Stream Vulnerability”—motion, mathematics, amblyopia, actions, and attention. J Vis 17:26.

2. Batschelet E (1981) Circular statistics in biology. New York, NY, US: Academic Press.

3. Bizley JK, Nodal FR, Bajo VM, Nelken I, King AJ (2007) Physiological and Anatomical Evidence for Multisensory Interactions in Auditory Cortex. Cereb Cortex 17:2172–2189.

4. Bollmann JH (2019) The Zebrafish Visual System: From Circuits to Behavior. Annu Rev Vis Sci 5:269–293.

5. Born RT, Bradley D (2005) Structure and Function of Visual Area MT. Annu Rev Neurosci 28:157–189.

6. Borst A, Groschner LN (2023) How Flies See Motion. Annu Rev Neurosci 46:17–37.

7. Bourne JA, Cichy RM, Kiorpes L, Morrone MC, Arcaro MJ, Nielsen KJ (2024) Development of Higher-Level Vision: A Network Perspective. J Neurosci 44:e1291242024.

8. Bourne JA, Rosa MGP (2006) Hierarchical development of the primate visual cortex, as revealed by neurofilament immunoreactivity: early maturation of the middle temporal area (MT). Cereb Cortex 16:405–414.

9. Brainard DH (1997) The Psychophysics Toolbox. Spat Vis 10:433–436.

10. Chang JT, Fitzpatrick D (2022) Development of visual response selectivity in cortical GABAergic interneurons. Nat Commun 13:3791.

11. Chaplin TA, Rosa MGP, Lui LL (2018) Auditory and Visual Motion Processing and Integration in the Primate Cerebral Cortex. Front Neural Circuits 12.

12. Chapman B, Stryker MP (1993) Development of orientation selectivity in ferret visual cortex and effects of deprivation. J Neurosci 13:5251.

13. Chapman B, Stryker MP, Bonhoeffer T (1996) Development of orientation preference maps in ferret primary visual cortex. J Neurosci 16:6443–6453.

14. Clemens JM, Ritter NJ, Roy A, Miller JM, Van Hooser SD (2012) The Laminar Development of Direction Selectivity in Ferret Visual Cortex. J Neurosci 32:18177–18185.

15. De Valois RL, Yund EW, Hepler N (1982) The orientation and direction selectivity of cells in macaque visual cortex. Vision Res 22:531–544.

16. Du J, Blanche TJ, Harrison RR, Lester HA, Masmanidis SC (2011) Multiplexed, High Density Electrophysiology with Nanofabricated Neural Probes. PLOS ONE 6:e26204.

17. Ellaway PH (1978) Cumulative sum technique and its application to the analysis of peristimulus time histograms. Electroencephalogr Clin Neurophysiol 45:302–304.

18. El-Shamayleh Y, Kiorpes L, Kohn A, Movshon JA (2010) Visual Motion Processing by Neurons in Area MT of Macaque Monkeys with Experimental Amblyopia. J Neurosci 30:12198–12209.

19. Gao WJ, Newman DE, Wormington AB, Pallas SL (1999) Development of inhibitory circuitry in visual and auditory cortex of postnatal ferrets: immunocytochemical localization of GABAergic neurons. J Comp Neurol 409:261–273.

20. Gao WJ, Wormington AB, Newman DE, Pallas SL (2000) Development of inhibitory circuitry in visual and auditory cortex of postnatal ferrets: Immunocytochemical localization of calbindin- and parvalbumin-containing neurons. J Comp Neurol 422:140–157.

21. Grootel TJV, Raghavan RT, Kelly JG, Movshon JA, Kiorpes L (2024) Responses to visual motion of neurons in the extrastriate visual cortex of macaque monkeys with experimental amblyopia. :2024.07.01.601564.

22. Homman-Ludiye J, Manger P, Bourne JA (2010) Immunohistochemical parcellation of the ferret (Mustela putorius) visual cortex reveals substantial homology with the cat (Felis catus). J Comp Neurol 518:4439–4462.

23. Hubel DH, Wiesel TN (1968) Receptive fields and functional architecture of monkey striate cortex. J Physiol 195:215–243.

24. Huxlin KR, Martin T, Kelly K, Riley M, Friedman DI, Burgin WS, Hayhoe M (2009) Perceptual Relearning of Complex Visual Motion after V1 Damage in Humans. J Neurosci 29:3981–3991.

25. Jarosiewicz B, Schummers J, Malik WQ, Brown EN, Sur M (2012) Functional biases in visual cortex neurons with identified projections to higher cortical targets. Curr Biol 22:269–277.

26. Khalil R, Contreras-Ramirez V, Levitt JB (2018) Postnatal refinement of interareal feedforward projections in ferret visual cortex. Brain Struct Funct 223:2303–2322.

27. Khalil R, Levitt JB (2014) Developmental remodeling of corticocortical feedback circuits in ferret visual cortex. J Comp Neurol 522:3208–3228.

28. Lempel AA, Nielsen KJ (2019) Ferrets as a Model for Higher-Level Visual Motion Processing. Curr Biol 29:179–191.

29. Lempel AA, Nielsen KJ (2021) Development of visual motion integration involves coordination of multiple cortical stages. eLife 10:e59798.

30. Li Y, Fitzpatrick D, White LE (2006) The development of direction selectivity in ferret visual cortex requires early visual experience. Nat Neurosci 9:676–681.

31. Li Y, Van Hooser SD, Mazurek M, White LE, Fitzpatrick D (2008) Experience with moving visual stimuli drives the early development of cortical direction selectivity. Nature 456:952–956.

32. Livingstone MS, Pack CC, Born RT (2001) Two-Dimensional Substructure of MT Receptive Fields. Neuron 30:781–793.

33. Mazurek M, Kager M, Van Hooser SD (2014) Robust quantification of orientation selectivity and direction selectivity. Front Neural Circuits 8.

34. Movshon JA, Rust NC, Kohn L, Kiorpes L, Hawken MJ (2004) Receptive-field properties of MT neurons in infant macaques. Perception 33:27.

35. Mulholland HN, Hein B, Kaschube M, Smith GB (2021) Tightly coupled inhibitory and excitatory functional networks in the developing primary visual cortex Feller MB, Moore T, Goodhill GJ, eds. eLife 10:e72456.

36. Mundinano I-C, Fox DM, Kwan WC, Vidaurre D, Teo L, Homman-Ludiye J, Goodale MA, Leopold DA, Bourne JA (2018) Transient visual pathway critical for normal development of primate grasping behavior. Proc Natl Acad Sci:201717016.

37. Murakami T, Matsui T, Ohki K (2017) Functional Segregation and Development of Mouse Higher Visual Areas. J Neurosci 37:9424–9437.

38. Murakami T, Matsui T, Uemura M, Ohki K (2022) Modular strategy for development of the hierarchical visual network in mice. Nature 608:578–585.

39. Murphy AJ, Hasse JM, Briggs F (2020) Physiological characterization of a rare subpopulation of doublet-spiking neurons in the ferret lateral geniculate nucleus. J Neurophysiol 124:432–442.

40. Niell CM, Scanziani M (2021) How Cortical Circuits Implement Cortical Computations: Mouse Visual Cortex as a Model. Annu Rev Neurosci 44:517–546.

41. Orban G (2008) Higher Order Visual Processing in Macaque Extrastriate Cortex. Physiol Rev 88:59–89.

42. Pelli DG (1997) The VideoToolbox software for visual psychophysics: transforming numbers into movies. Spat Vis 10:437–442.

43. Philipp R, Distler C, Hoffmann K-P (2006) A motion-sensitive area in ferret extrastriate visual cortex: an analysis in pigmented and albino animals. Cereb Cortex 16:779–790.

44. Popovic M, Stacy AK, Kang M, Nanu R, Oettgen CE, Wise DL, Fiser J, Van Hooser SD (2018) Development of Cross-Orientation Suppression and Size Tuning and the Role of Experience. J Neurosci 38:2656–2670.

45. Price DJ, Zumbroich TJ, Blakemore C (1988) Development of Stimulus Selectivity and Functional Organization in the Suprasylvian Visual Cortex of the Cat. Proc R Soc Lond B Biol Sci 233:123–163.

46. Raiguel SE, Lagae L, Gulyàs B, Orban GA (1989) Response latencies of visual cells in macaque areas V1, V2 and V5. Brain Res 493:155–159.

47. Raiguel SE, Xiao D-K, Marcar VL, Orban GA (1999) Response Latency of Macaque Area MT/V5 Neurons and Its Relationship to Stimulus Parameters. J Neurophysiol 82:1944–1956.

48. Ritter NJ, Anderson NM, Van Hooser SD (2017) Visual Stimulus Speed Does Not Influence the Rapid Emergence of Direction Selectivity in Ferret Visual Cortex. J Neurosci 37:1557–1567.

49. Roy A, Wang S, Meschede-Krasa B, Breffle J, Van Hooser SD (2020) An early phase of instructive plasticity before the typical onset of sensory experience. Nat Commun 11:11.

50. Rust NC, Mante V, Simoncelli EP, Movshon JA (2006) How MT cells analyze the motion of visual patterns. Nat Neurosci 9:1421–1431.

51. Schmolesky MT, Wang Y, Hanes DP, Thompson KG, Leutgeb S, Schall JD, Leventhal AG (1998) Signal Timing Across the Macaque Visual System. J Neurophysiol 79:3272–3278.

52. Smith GB, Sederberg A, Elyada YM, Van Hooser SD, Kaschube M, Fitzpatrick D (2015) The development of cortical circuits for motion discrimination. Nat Neurosci 18:252–261.

53. Smith IT, Townsend LB, Huh R, Zhu H, Smith SL (2017) Stream-dependent development of higher visual cortical areas. Nat Neurosci 20:200–208.

54. Spear PD, Tong L, McCall MA, Pasternak T (1985) Developmentally induced loss of direction-selective neurons in the cat’s lateral suprasylvian visual cortex. Dev Brain Res 20:281–285.

55. Stacy AK, Schneider NA, Gilman NK, Van Hooser SD (2023) Impact of Acute Visual Experience on Development of LGN Receptive Fields in the Ferret. J Neurosci 43:3495–3508.

56. Thiele A, Distler C, Korbmacher H, Hoffmann K-P (2004) Contribution of inhibitory mechanisms to direction selectivity and response normalization in macaque middle temporal area. Proc Natl Acad Sci 101:9810–9815.

57. Thiele A, Herrero JL, Distler C, Hoffmann K-P (2012) Contribution of Cholinergic and GABAergic Mechanisms to Direction Tuning, Discriminability, Response Reliability, and Neuronal Rate Correlations in Macaque Middle Temporal Area. J Neurosci 32:16602–16615.

58. Van Hooser SD, Li Y, Christensson M, Smith GB, White LE, Fitzpatrick D (2012) Initial neighborhood biases and the quality of motion stimulation jointly influence the rapid emergence of direction preference in visual cortex. J Neurosci 32:7258–7266.

59. Vaney DI, Sivyer B, Taylor WR (2012) Direction selectivity in the retina: symmetry and asymmetry in structure and function. Nat Rev Neurosci 13:194–208.

60. Warner CE, Kwan WC, Bourne JA (2012) The Early Maturation of Visual Cortical Area MT is Dependent on Input from the Retinorecipient Medial Portion of the Inferior Pulvinar. J Neurosci 32:17073–17085.

61. Wei W, Feller MB (2011) Organization and development of direction-selective circuits in the retina. Trends Neurosci 34:638–645.

62. Weiskrantz L (1996) Blindsight revisited. Curr Opin Neurobiol 6:215–220.

63. Weiskrantz L, Barbur JL, Sahraie A (1995) Parameters affecting conscious versus unconscious visual discrimination with damage to the visual cortex (V1). Proc Natl Acad Sci 92:6122–6126.

64. Weliky M, Bosking W, Fitzpatrick D (1996) A systematic map of direction preference in primary visual cortex. Nature 379:725–728.

65. Zhang B, Smith EL, Chino YM (2008) Postnatal development of onset transient responses in macaque V1 and V2 neurons. J Neurophysiol 100:1476–1487.

