## Supplemental Figures for "Early development of direction selectivity in higher visual cortex"

Supplement Figure 1: Stimulus parameters across areas and age groups

(A) Stimulus spatial frequency as a function of age and recording location. Each symbol indicates one experiment included in the data set. Note that a small random jitter is added to the spatial frequency values for plotting purposes only so that all experiments remain visible in the plot. Dashed lines indicate values that were used in at least one experiment. Symbol color and shape indicate experiments with simultaneous recordings in V1 and PMLS, and experiments in which only PMLS or V1 data was recorded (see legend in (C)).

(B) Temporal frequency as a function of age and recording location, using the same plot format as in (A).

(C) Inter-stimulus interval values across experiments (same format as (A)).

Supplement Figure 2: Development of DS and OS, quantified using DSI and OSI.

(Supplement to Fig. 1 & 2)

(A) Average DSI value per animal, plotted as a function of the animal's age. Same format as Fig. 1D otherwise.

(B, C) Cumulative DSI distributions for SU data recorded in PMLS and V1, divided into 3 age groups. Same format as Fig. 1E & F.

(D) Average OSI per-animal as a function of age; same format as Fig. 1D.

(E, F) Cumulative OSI distributions for SU data recorded in PMLS and V1, divided into 3 age groups. Same format as Fig. 1E & F.

(G, H) Cumulative distributions of null responses relative to preferred responses for both areas and all age groups.

(I) Mean null/preferred responses for PMLS and V1 as a function of age. Data were binned into the same age windows used for Fig. 1G. No normalization was used here, in contrast to the data in Fig. 1G. Error bars: SEM.

See Table 13 for full statistical results.

Supplement Figure 3: Per-animal results for the training cohorts.

(Supplement to Fig. 4)

(A) Effect of exposure to different stimuli for $L_{dir}$ levels in V1 (left: drifting gratings, middle: gray screen only, right: flashing gratings). In each plot, each column of data points reflects data collected before and after training in one animal, with every data point corresponding to the $L_{dir}$ value of a single visually responsive MU site. The color code indicates corresponding pre- and post-training data for the same animal for a single plot only; it does not indicate correspondence between plots, including V1 and PMLS data for the same training stimulus (i.e. animal 1 for V1 does not have to correspond to animal 1 for PMLS). The black line indicates the before and after $L_{dir}$ levels across animals as estimated from the fixed effect in the LME model. Red lines are per-animal $L_{dir}$ levels as estimated by both the fixed effect and random, per animal effects.

(B) Effect of different training stimuli on $L_{dir}$ levels in PMLS. Same format as in (A).

(C) Effect of different training stimuli on $L_{ori}$ levels in V1. Same format as in (A).

(D) Effect of different training stimuli on $L_{ori}$ levels in PMLS. Same format as in (A).

See Tables 9 & 10 for full statistics.

Supplement Figure 4: Comparison of responses evoked by flashing and drifting gratings.
(Supplement to Fig. 4)
(A) Correlation in $L_{ori}$ training effect size between V1 and PMLS. Each symbol corresponds to data from a single animal for which both V1 and PMLS data was available, with training effect size determined by the LME model. Error bars: SE estimated for the per-animal random slope. Different training cohorts are indicated by different symbols and colors.
(B) Impact of pre-training $L_{dir}$ levels on training effect size for V1 for all 3 training cohorts. Pre-training levels and training effect sizes were determined from the LME model as before (see legend for the different groups).
(C) Same as (B), but for $L_{ori}$ levels in V1.
(D) Impact of pre-training $L_{dir}$ levels on training effect size in PMLS for the 3 cohorts. Same format as (B).
(E) Same as (D), but for PMLS $L_{ori}$ levels.

Supplement Figure 5: Comparison of responses evoked by flashing and drifting gratings.
(Supplement to Fig. 4)
Cumulative distributions of peak rate, response duration and average rate for MU sites recorded in V1 and PMLS in response to drifting and flashing gratings (same color scheme as in Fig. 4I). The legend indicates the outcome of rank-sum tests between flashing and drifting gratings per area (***: $p<.001$). No tests were performed for the peak rate as the group-level ANOVA indicated no differences between the stimulus conditions (see Table 11).

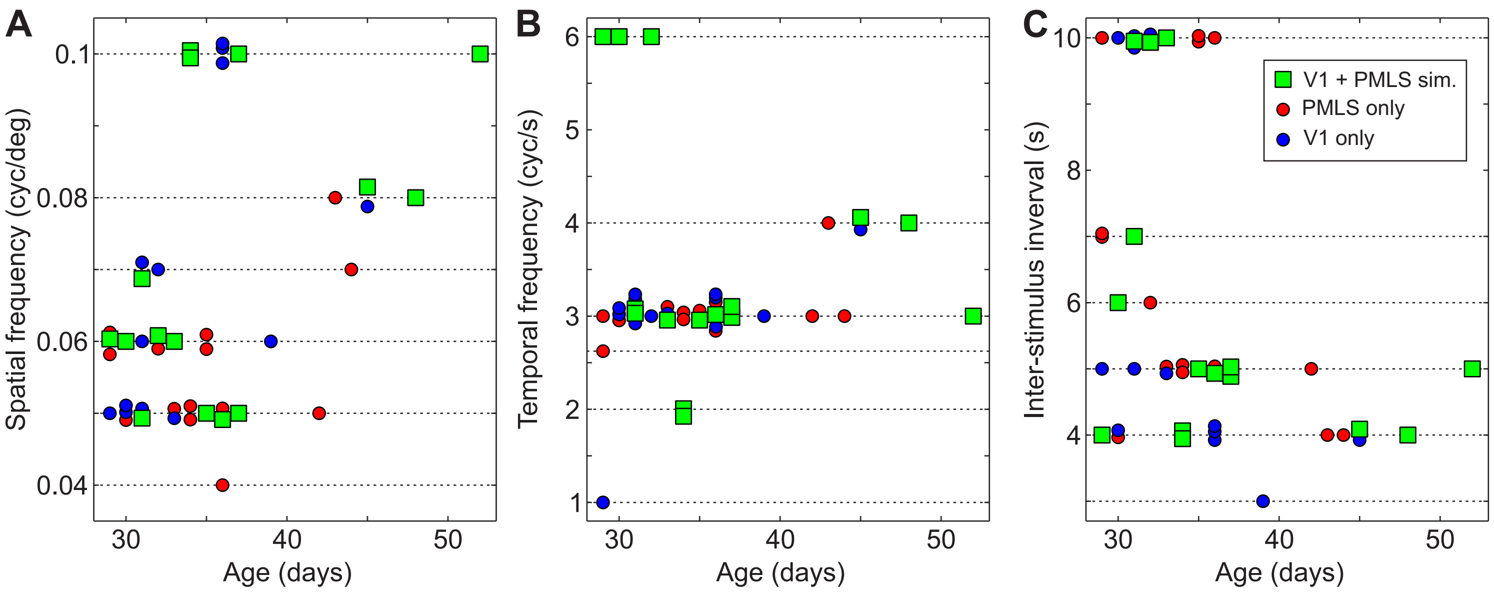


**Suppl. Figure 1**

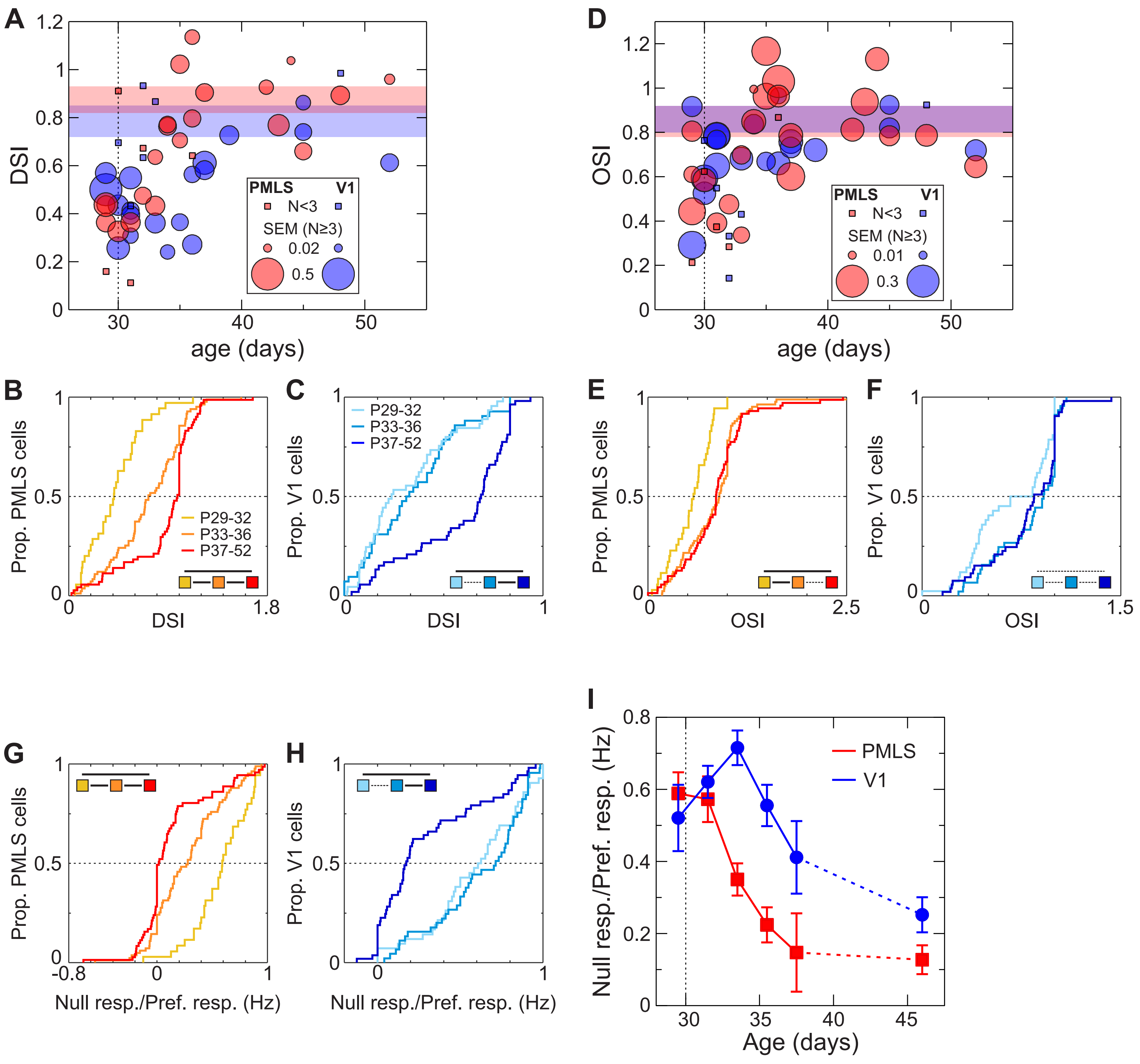


**Suppl. Figure 2**

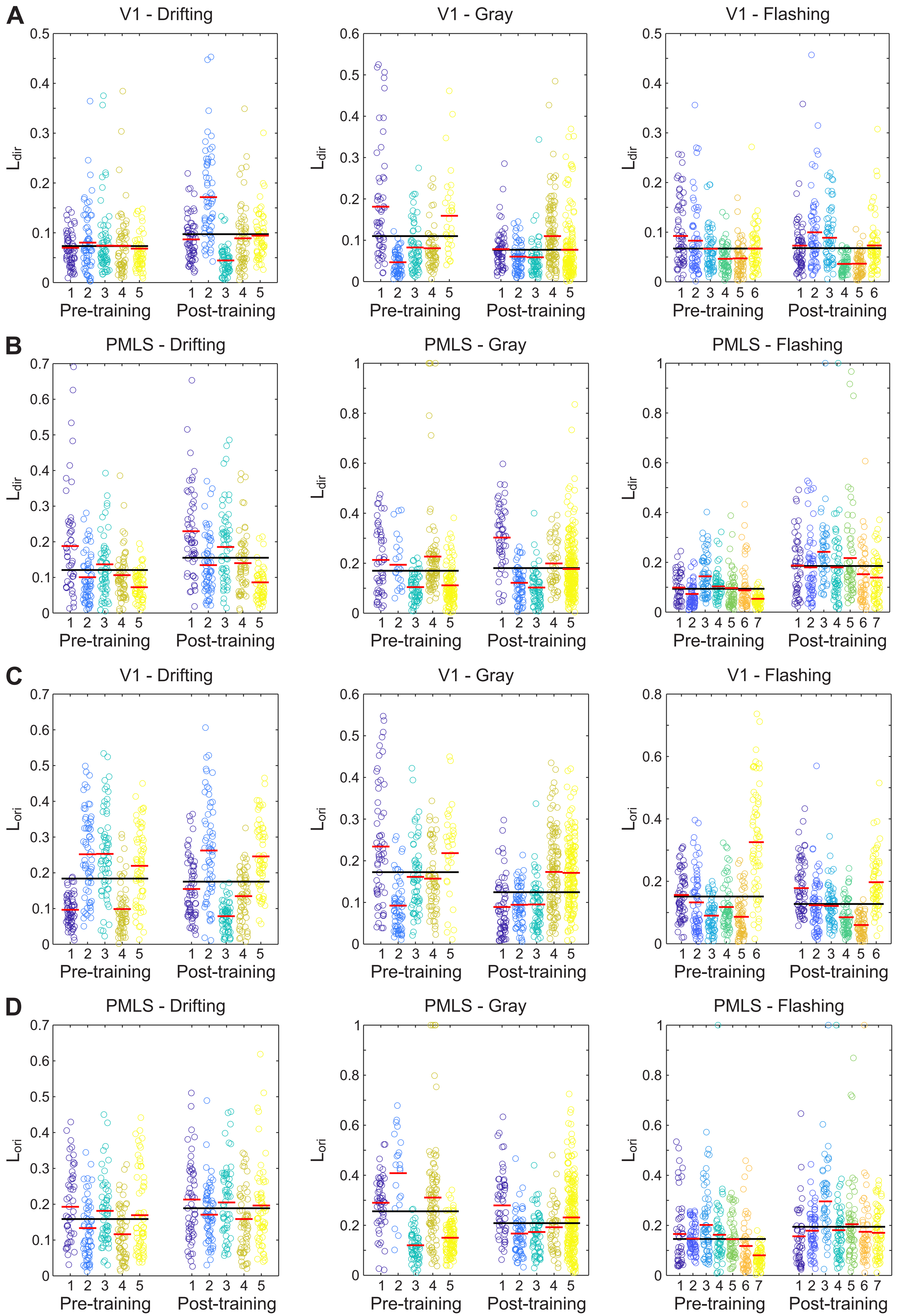


**Suppl. Figure 3**

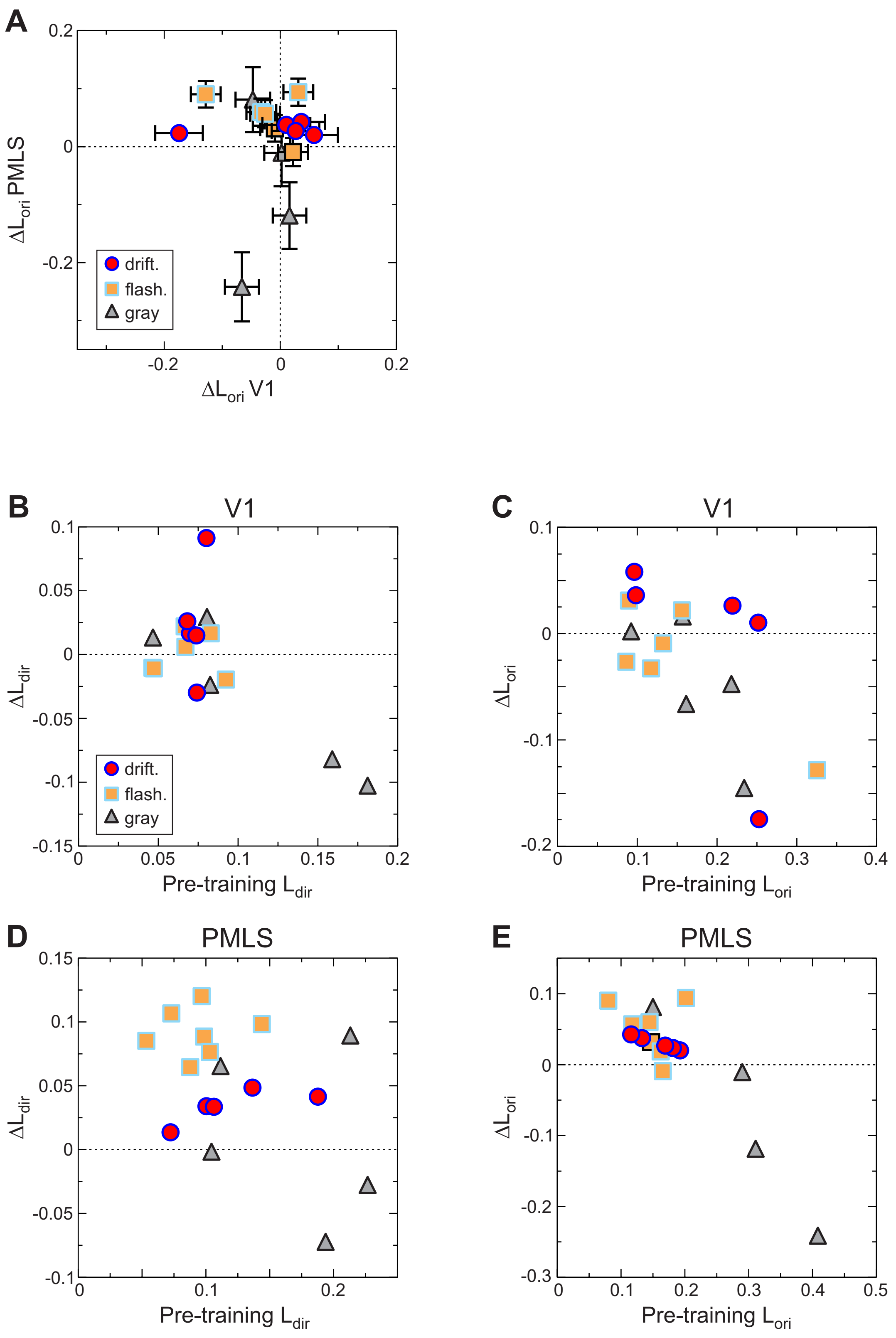


**Suppl. Figure 4**

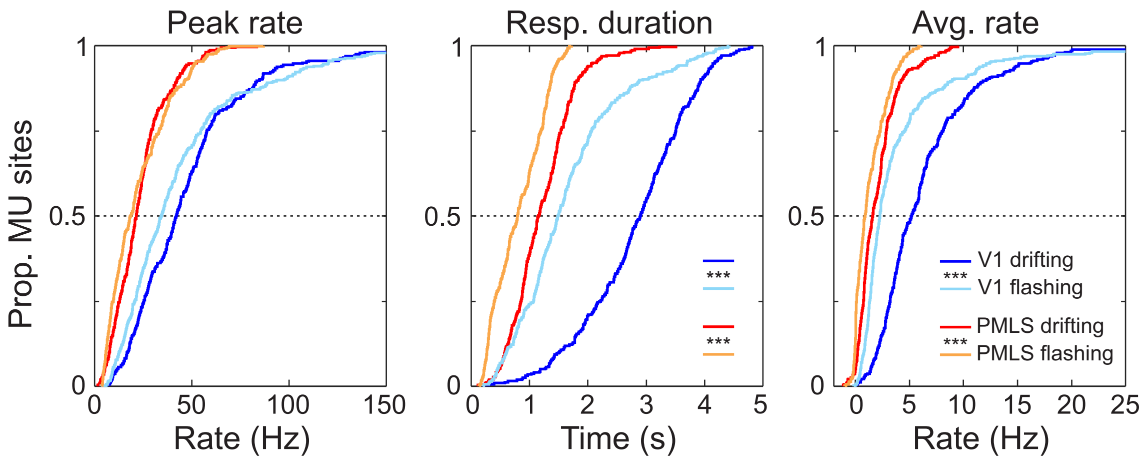


**Suppl. Figure 5**
